# Generative artificial intelligence GPT-4 accelerates knowledge mining and machine learning for synthetic biology

**DOI:** 10.1101/2023.06.14.544984

**Authors:** Zhengyang Xiao, Wenyu Li, Hannah Moon, Garrett W. Roell, Yixin Chen, Yinjie J. Tang

## Abstract

Knowledge mining from synthetic biology journal articles for machine learning (ML) applications is a labor-intensive process. The development of natural language processing (NLP) tools, such as GPT-4, can accelerate the extraction of published information related to microbial performance under complex strain engineering and bioreactor conditions. As a proof of concept, we used GPT-4 to extract knowledge from 176 publications on two oleaginous yeasts (*Yarrowia lipolytica* and *Rhodosporidium toruloides*). After integration with a molecule inventory database, the outcome is a total of 2037 data instances and 28 features, which serve as machine learning inputs. The structured datasets enabled ML approaches (e.g., a random forest model) to predict Yarrowia fermentation titers with high accuracy (R^2^ of 0.86 for unseen test data). Via transfer learning, the trained model could also assess the production capability of the non-conventional yeast, *R. toruloides*, for which there are fewer published reports. This work demonstrated the potential of generative artificial intelligence to speed up information extraction from research articles, thereby improving design-build-test-learn (DBTL) cycles for commercial biomanufacturing development.

## Introduction

Synthetic biology tools can engineer microbes for sustainable biomanufacturing. To develop microbial workhorses, researchers rely on trial and error for breakthroughs due to the complex nature of biological systems. Model predictions of cell performance are key to reduce experimental trials and improve strain development effectiveness. However, mechanistic models (such as genome scale model) have difficulties in incorporating all influential factors to simulate microbial production^1^. On the other hand, data driven models (i.e., ML) have been applied to predict fermentation titers^2–4^, optimize bioprocesses^5–7^, and recommend strain engineering approaches^8, 9^. The drawback of ML is that it requires large sets of experimental data for training. Therefore, knowledge mining from published journal articles can be an inexpensive strategy to train ML models. However, manually extracting data from a large number of articles is labor-intensive and prone to human errors and inconsistencies in quality, because reported data often lack a standardized format^10, 11^ and substantial efforts are needed to interpret information and organize it into ML-ready data^12^.

NLP, a branch of AI, can process text at a large scale, enabling topic organization in published articles^13^. It has also been utilized to track adverse drug events from electronic health record notes^14^. A recent tipping point in the field of NLP was the release of GPT-4^15^, which shows ‘sparks’ of artificial general intelligence^16^ to rapidly parse text based on user-provided context^15^. Leveraging GPT-4, relevant bioprocess features and outcomes from published papers can be extracted for rapid database growth. Since GPT-4 is unable to offer quantitative predictions of microbial productions (**Supplementary Figure 1**), particularly for non-model species, this study aims to integrate GPT-4 with ML algorithms to predict the fermentation titer from various microbial cell factories ^2, 17^. As a proof of concept, GPT-4 is used to extract knowledge from articles on an industrial yeast *Yarrowia lipolytica*. After human supervision, these published case studies can be transformed into data samples (also called instances). Each instance includes both outputs (product type and titer) and inputs (i.e., numerical or categorical features). These feature variables include bioprocess conditions (e.g., medium composition, substrate types, and bioreactor types) and metabolic pathway information (e.g., the number of enzyme steps for product synthesis). All data samples have been uploaded to a database (impact-database.com) that can be used to train ML models. Moreover, *Rhodosporidium toruloides* is a novel yeast that has recently gained research attention for its high lipid content^18, 19^ and native carotenoid production^20, 21^. But the literature on *Rhodosporidium* is sparse^22^. Here, we demonstrate that transfer learning (TL) can transfer knowledge from well-studied domains (*Yarrowia*-trained model) to understand less-studied scenarios, reducing computational costs and speeding up the learning process^23, 24^. In summary, our study highlights the promising role of NLP for more efficient data collection and rapid centralization of published synthetic biology studies. For the first time, we demonstrate the integration of GPT with knowledge engineering and ML for predicting different microbial cell factories. The lessons from this study will also improve human supervision and prompt engineering for future GPT applications. A standardized workflow for GPT-4 data extraction and future applications is shown in **Figure 1**.

**Figure 1.**
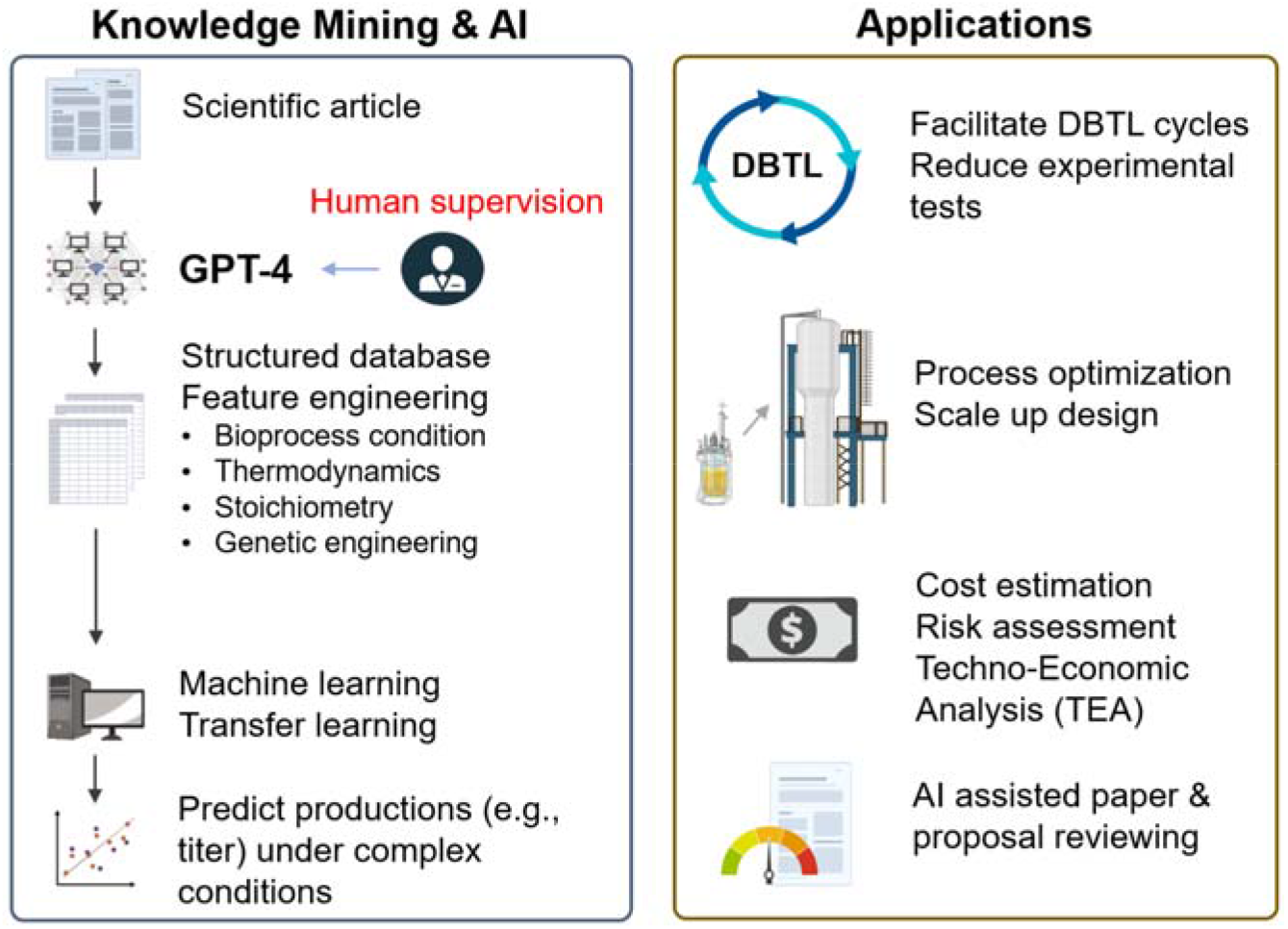
GPT-4 knowledge mining for ML (Left) and potential AI applications. (assist biomanufacturing design, commercial decision, or project quality/risk assessment).

## Results and discussion

### Extraction of ML features and datasets from synthetic biology papers via GPT-4 workflow

ML approaches require a large amount of experimental data to correlate ML inputs (features) to outputs (productions). Since biomanufacturing literature presents a wealth of strain construction and bioprocess engineering case studies, the construction of a database from published papers may broadly support ML applications. Previous database construction efforts, like LASER^10^, have stored knowledge in ways that are useful for metabolic engineers, but these databases have not been organized or transformed for direct ML applications. On the other hand, this study involves knowledge mining and feature engineering, which aims to filter erroneous/redundant information and capture the features that independently affect bio-production. In this study, biomanufacturing features are based on bioreactor conditions and genetic engineering methods. Manual extraction of these features and production outcomes can be time-consuming, so GPT-4 is used to overcome this challenge. The general principle is dividing complex tasks into small deliverables. Due to GPT-4’s max context window of 8,192 tokens, the sections of each scientific article, including abstract, materials and methods, results, and data tables, were manually separated into text files. Prompts were then added to the beginning of each section depending on their contents (**Table 1**). Then GPT-4 summarizes the information from experiment results and methods into accessible tables (See examples in **Supplementary File 1**).

**Table 1.**
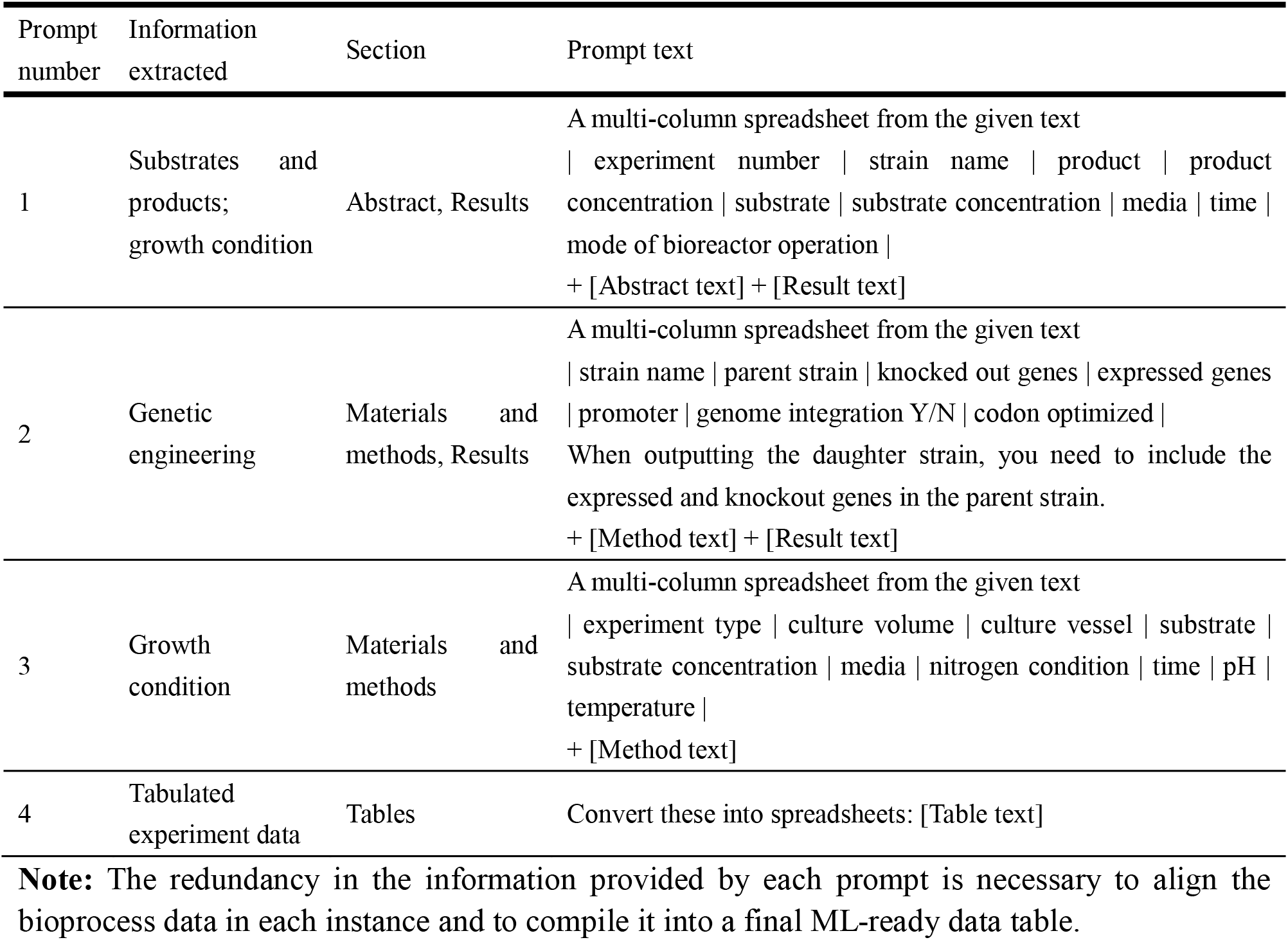
Prompts (questions for GPT) used for GPT-4 data extraction.

### Quality tests for data extraction by ChatGPT

Data extraction without losing important knowledge is challenging because data reporting from publications is often sparse and inconsistent. To test GPT applicability, we started to use GPT-3.5 API on March 15^th^, 2023 to extract *Rhodosporidium* fermentation data from journal articles. The output of one PhD student using the GPT-enhanced workflow for one week is illustrated in Figure 2a. When GPT-3.5 was used, on average 11.7 papers were extracted per day (8 work hours). After the release of GPT-4 on 3/15, 25 papers were extracted on 3/16. We tested the correctness of data extracted by GPT-3.5 in 10 papers. Compared with a previous manually extracted dataset^2^, we found that titer data extracted by GPT-3.5 were 74% correct. Some of the erroneous data was obvious as they included numbers that were consecutive, repeated, or not present in the article. With minor user discretion to fix the outputs, the extracted titer data was found to be 89% correct. When GPT-4 extracted data from the test set of 10 *Yarrowia* papers, we found no errors in titer data. We summarized the data location, output format, and correctness in (**Supplemental Table 2**). In total, 366 data instances were obtained from 60 *Rhodosporidium* articles.

**Figure 2.**
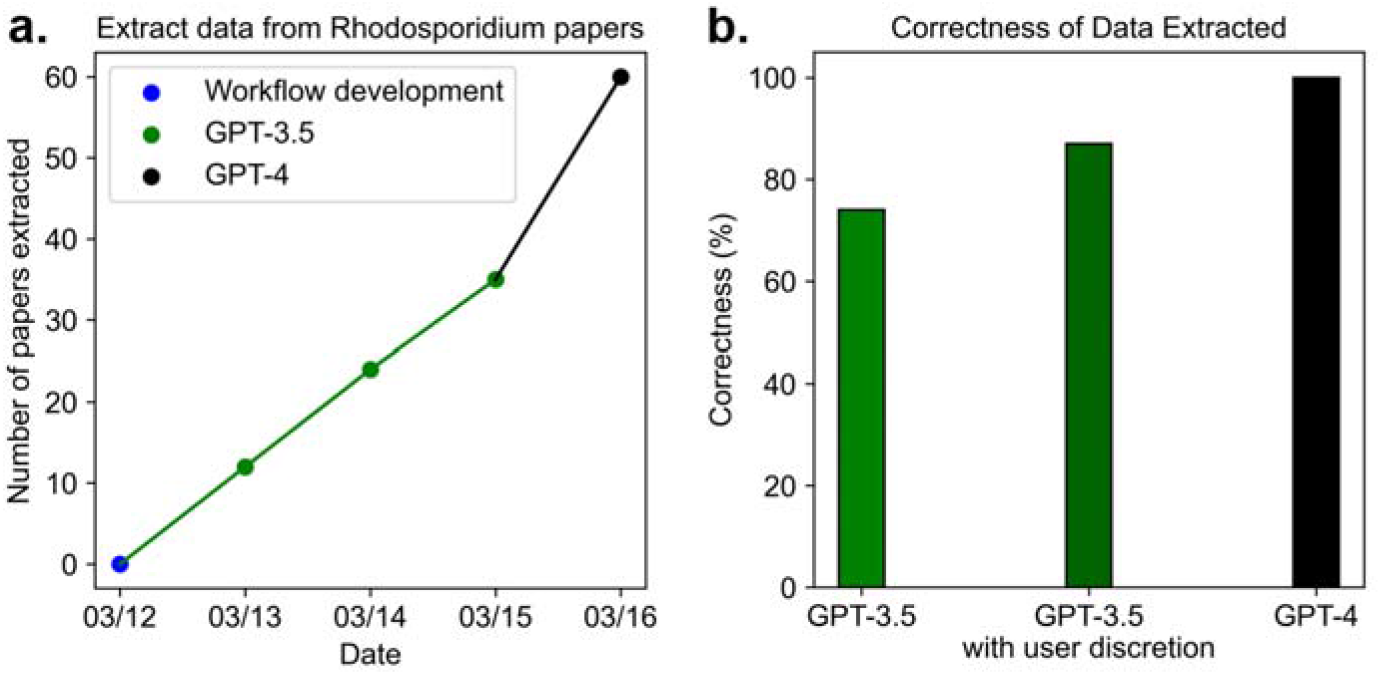
Speed and correctness difference between GPT-3.5 and GPT-4. **a.** Number of *Rhodosporidium* papers processed in five days by a single user. **b.** The correctness of data extracted from a test set of 10 *Yarrowia* articles that contained 115 fermentation instances. Note: all data were manually inspected for error before use in machine learning.

At this stage, human supervision is still a necessity to ensure GPT output’s accuracy. To extract data from one paper, the workflow typically took 20 minutes: labeling and dividing article sections ➔ entering prompts on ChatGPT website ➔ recording GPT response ➔ combining the results into ML-ready dataset ➔ quality check. This workflow is an improvement over manual reading because: 1) it does not rely on the expertise of a single person and can be parallelized across a team, 2) it does not require high levels of effort for data recording, and 3) it extracts data reproducibly, and the extracted data can be checked for errors in a *targeted* manner rather than laboriously reading each section^25^, and 4) it is adaptable to automation once the GPT-4’s Application Programming Interface (API) is available.

### Case study: GPT-4 assisted database construction for *Yarrowia lipolytica* biomanufacturing

*Yarrowia lipolytica* is an industrially important yeast for bio-productions^26^. Our previous study has extracted information manually from ∼100 *Yarrowia* papers (∼3000 instances)^2^, which took an experienced graduate student over 400 working hours. Via the GTP-4 workflow, ∼1670 additional data instances from 115 papers were extracted and organized into 28 features that may influence production titer (**Table 2**). Apart from the experiment data extracted by GPT-4 from texts, we also developed a ***molecule inventory*** to include data such as thermodynamics properties, biological production pathway steps, precursors, and cofactor costs. This molecule inventory is an important part of our ImpactDB online database, and we will continuously update it in the future. With this centralized inventory, we can directly search for the information for each substrate/product and fill in ML features, saving precious time during database construction.

**Table 2.**
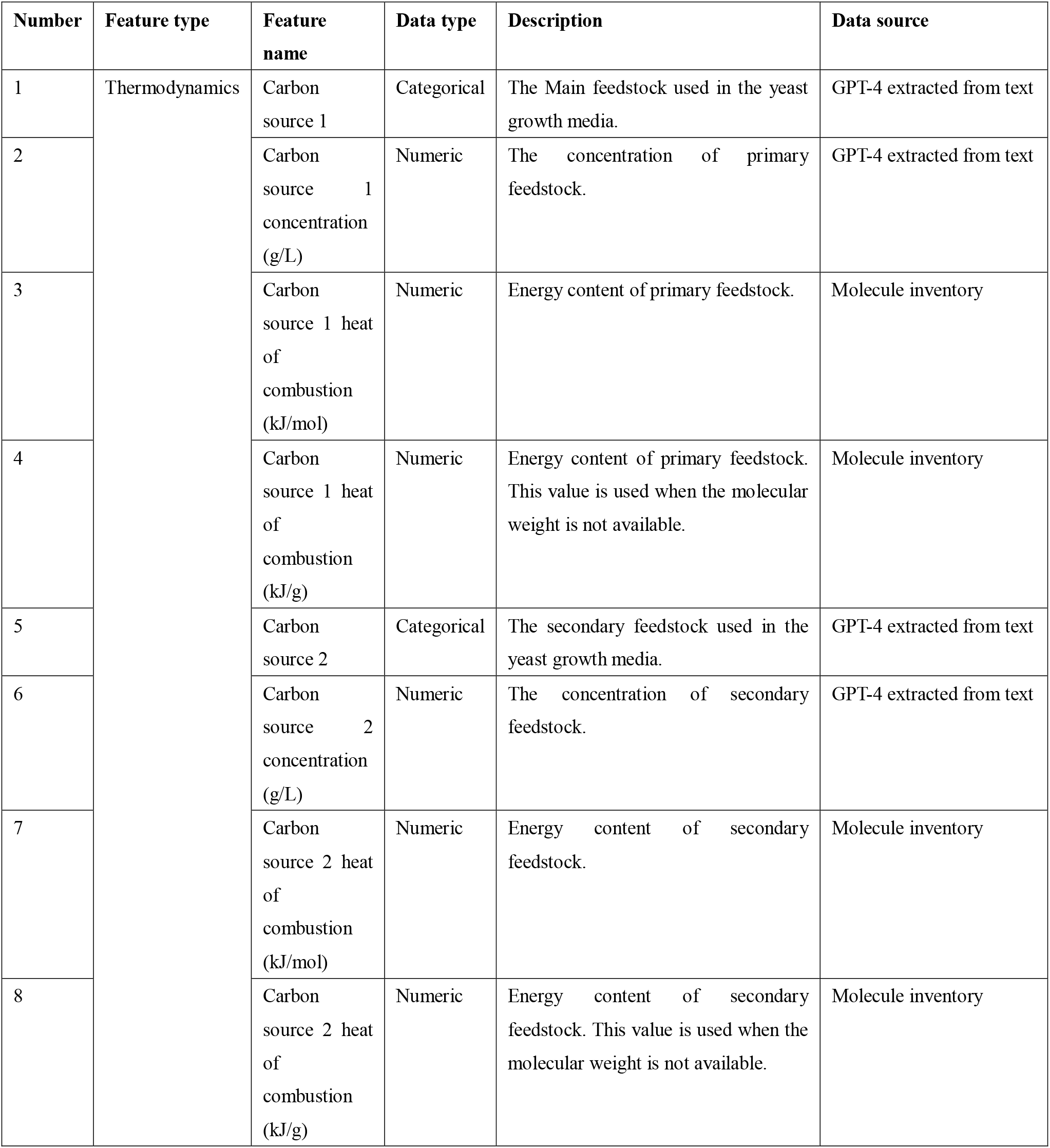
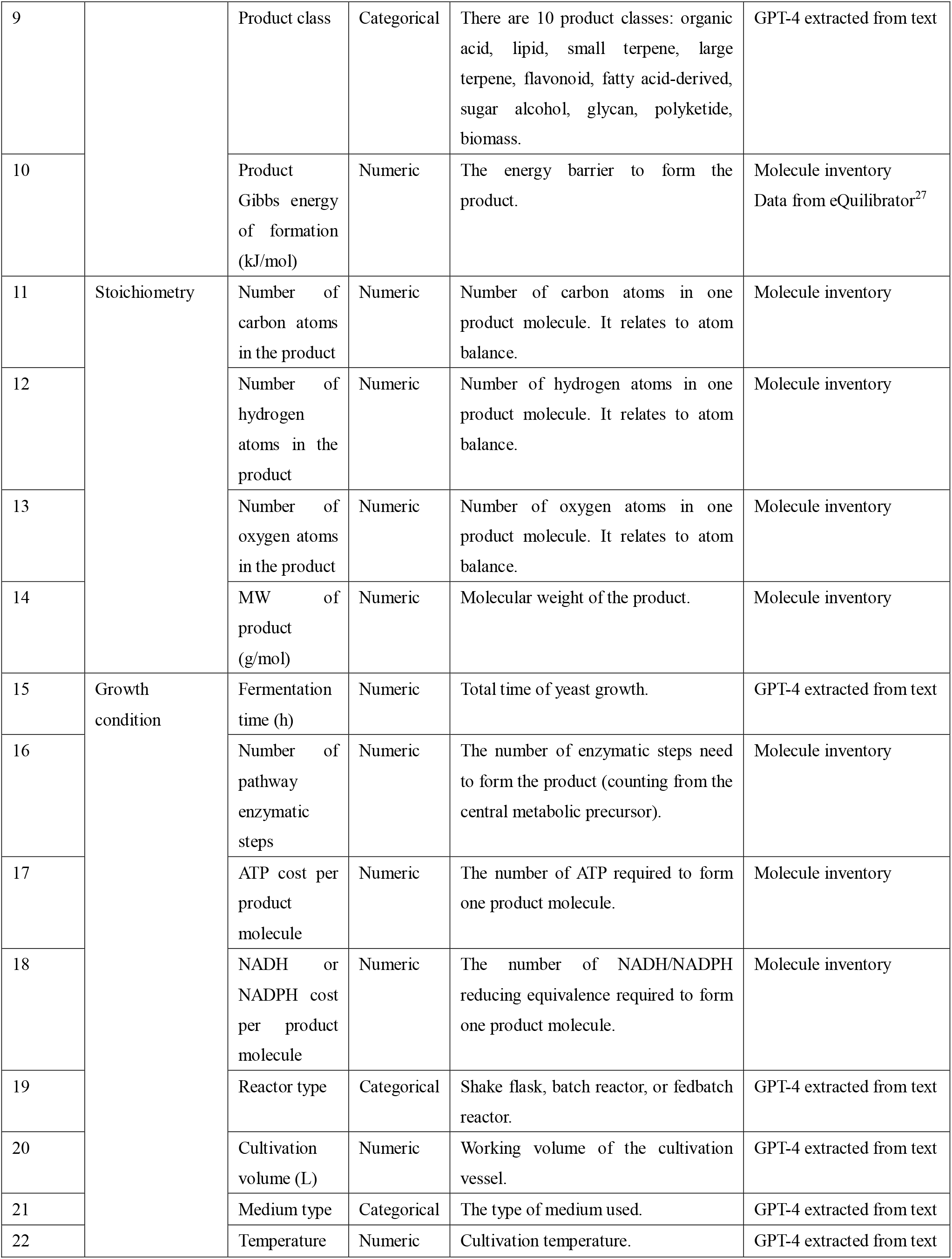
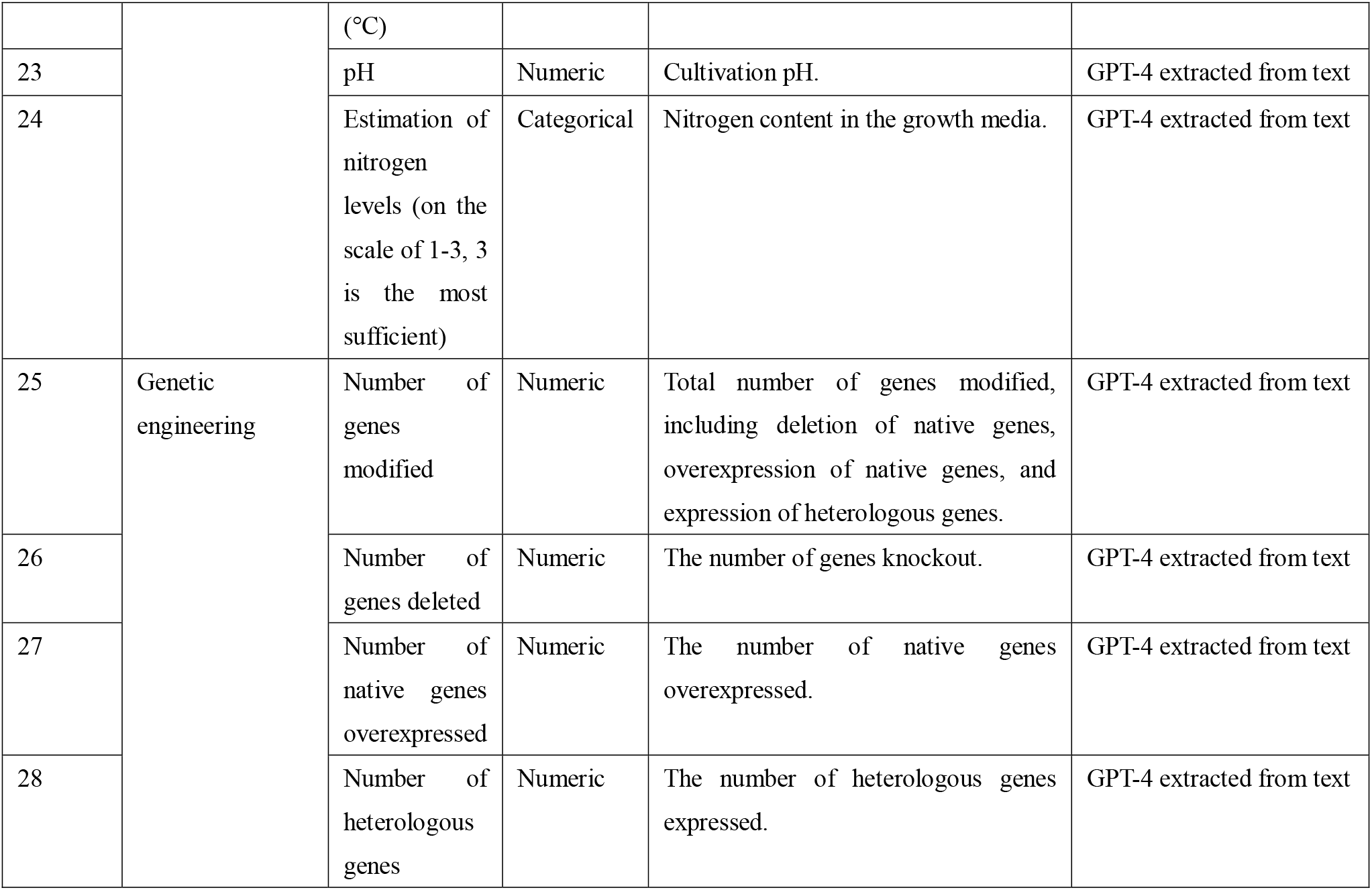
Features used in machine learning. (More details can be found in impact-database.com)

For further validating the applicability of GPT-4, the manually extracted data was compared with GPT extracted data by computing feature importance, feature variances, and principal component analysis (PCA). The GPT extracted data had a similar distribution of feature importance to that of the manually extracted data (**Figure 3a**), suggesting that the newly generated data followed similar patterns to the manually extracted data. Interestingly, the GPT datasets had higher feature variance than the manually extracted dataset for 19 of 28 features (**Figure 3b**). Moreover, a principal component analysis (PCA) showed that with K-means converged to the optimal solution and the same silhouette score, the data extracted by GPT showed 7% higher mean distance between clusters (**Figure 4**). The PCA loadings further indicated that the clustering of the manually extracted dataset was governed mainly by carbon source and product cofactor cost (**Supplementary Figure 2**). In contrast, GPT extracted data was clustered according to culture condition and genetic engineering features, in addition to carbon source and cofactor cost. These findings suggest that GPT-4 can capture more distinctiveness within papers and reason through complex contextual data to generate less biased biomanufacturing instances.

**Figure 3.**
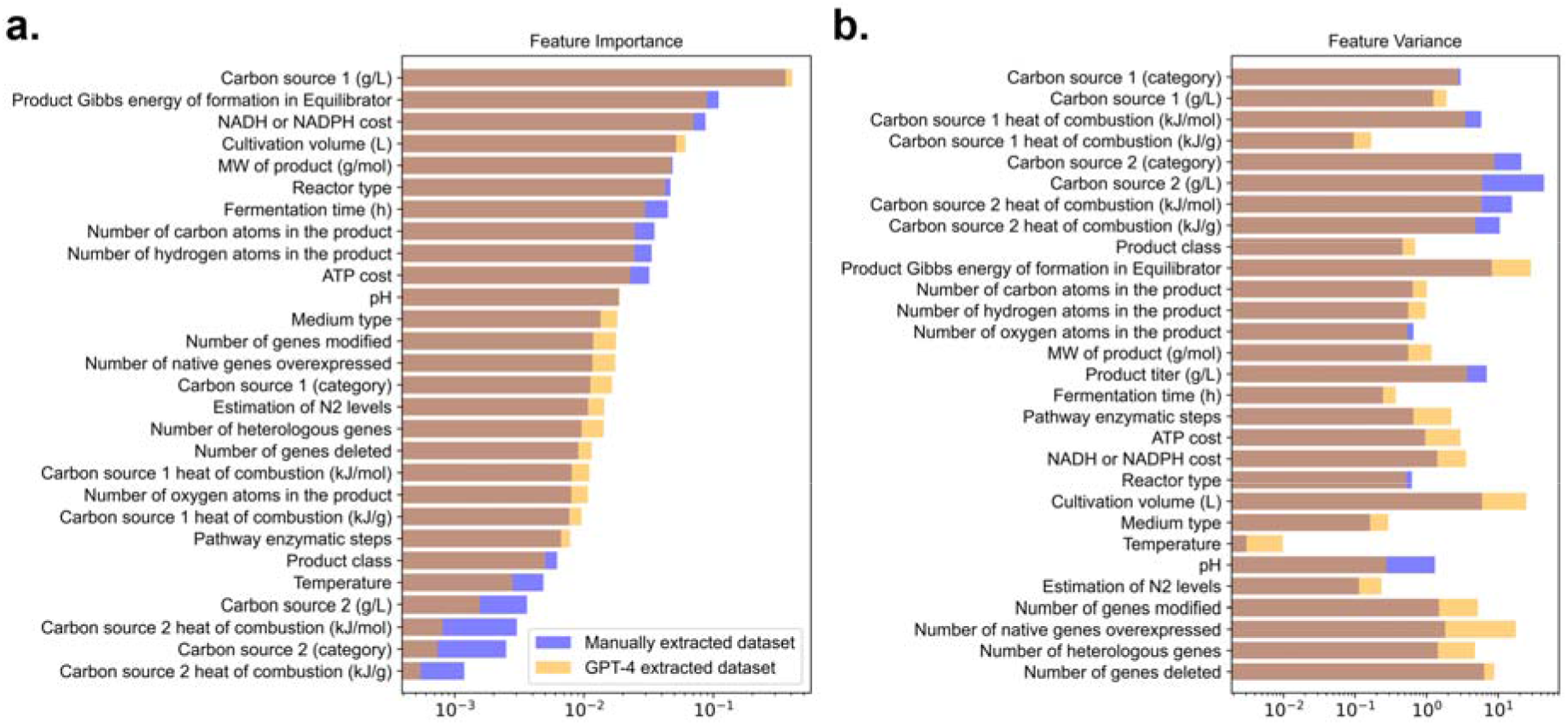
Comparison of the manually extracted *Yarrowia* dataset with the GPT-4 extracted *Yarrowia* dataset. **a.** Feature importance as determined using a random forest regressor, ranked from high to low. **b.** Normalized feature variance. Legend: Purple is the manually extracted dataset, and yellow is GPT-4 extracted dataset.

**Figure 4.**
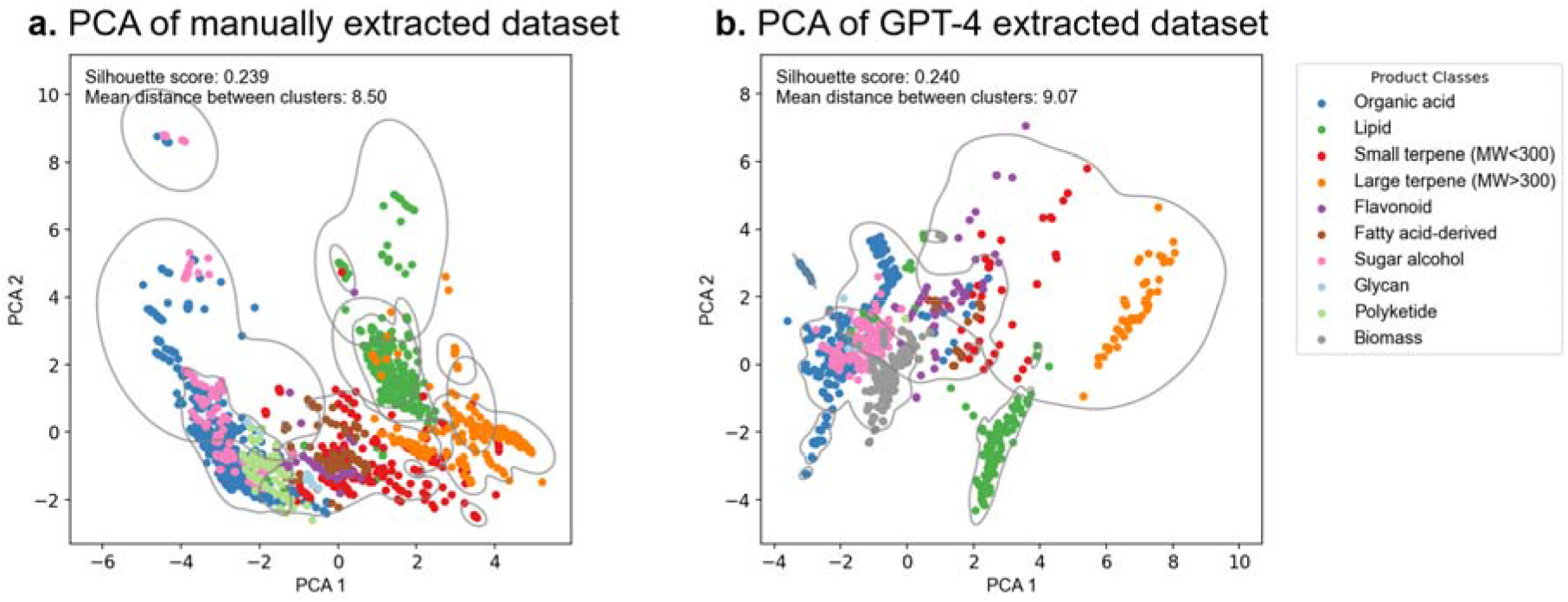
PCA using K-means unsupervised learning. **a)** PCA of the manually extracted dataset. **b)** PCA of GPT-4 extracted dataset. Note the axis scale difference between panels **a** and **b**.

### Leveraging the GPT-constructed database to predict *Y. lipolytica* fermentation titer

Fermentation titer determines the bioprocess economy and is the most important parameter for microbial cell factories. The GPT-assisted database construction can support the quantitative prediction of yeast fermentation titers under various conditions. Specifically, *Y. lipolytica* fermentation instances formed a comprehensive database to train ML models. We conducted a comparative test of seven classical ML algorithms (**Supplementary Figure 3**, support vector machine (SVM), gaussian process (GP), multi-layer perceptron (MLP), random forest (RF), extreme gradient boosting (XGBoost), k-nearest neighbors (KNN) and linear regression). Based on unseen testing data and ML prediction R^2^ (predicted titer vs. reported titer), ML methods including linear regression and linear SVM performed the least well, suggesting that the titer prediction cannot be represented accurately by a linear relationship. A fully connected 2-layer neural network did not give the best performance either. Overall, the RF model achieved the best accuracy: R^2^ of 0.86 on unseen test instances (**Figure 5a**). Therefore, we chose to use the RF ensemble learner to predict production titer, based on a combination of substrate concentrations, thermodynamics data, growth conditions, and engineered genes. After this point, we did not scale our train/test data, thus retaining their physical meaning. The test set performance was robust, with an average R^2^ of 0.80 across 50 random data splits. This represents a moderate improvement compared to our previous study^2^, which reported an average R^2^ of 0.76 in 10 random data splits (p≤0.005, student *t* test). The test set performance of the RF regressor was strong for nearly all product classes: organic acid, lipid, terpene, flavonoid, fatty acid-derived compound, sugar alcohol, glycan, and polyketide (**Figure 5b-k**). The new *Y. lipolytica* ML model, trained on a database approximately 50% larger than the previous model, also showed general improvement for titer predictions of small terpene and polyketide products. However, the new model still struggles to explain the data related to large terpenes, as indicated by the region with horizontal consecutive dots in the figure, where the same set of features results in a wide range of titers. This is because some key biomanufacturing features are still missing from knowledge mining. For instance, a recent study removed lycopene substrate inhibition by site-mutagenesis enzyme engineering, and it was able to achieve the highest β-carotene titer ever reported, 39.5 g/L^28^. However, the features in the collected database are not able to account for the substrate inhibition to production pathways. Also, our feature selection does not consider compartmentalization effects, as recent research showed that fermentation yields can be improved through cytoplasmic-peroxisomal engineering^29–31^. Additionally, the DNA sequence of key genes may be used as new ML inputs but still requires significant efforts for feature engineering^32, 33^.

**Figure 5.**
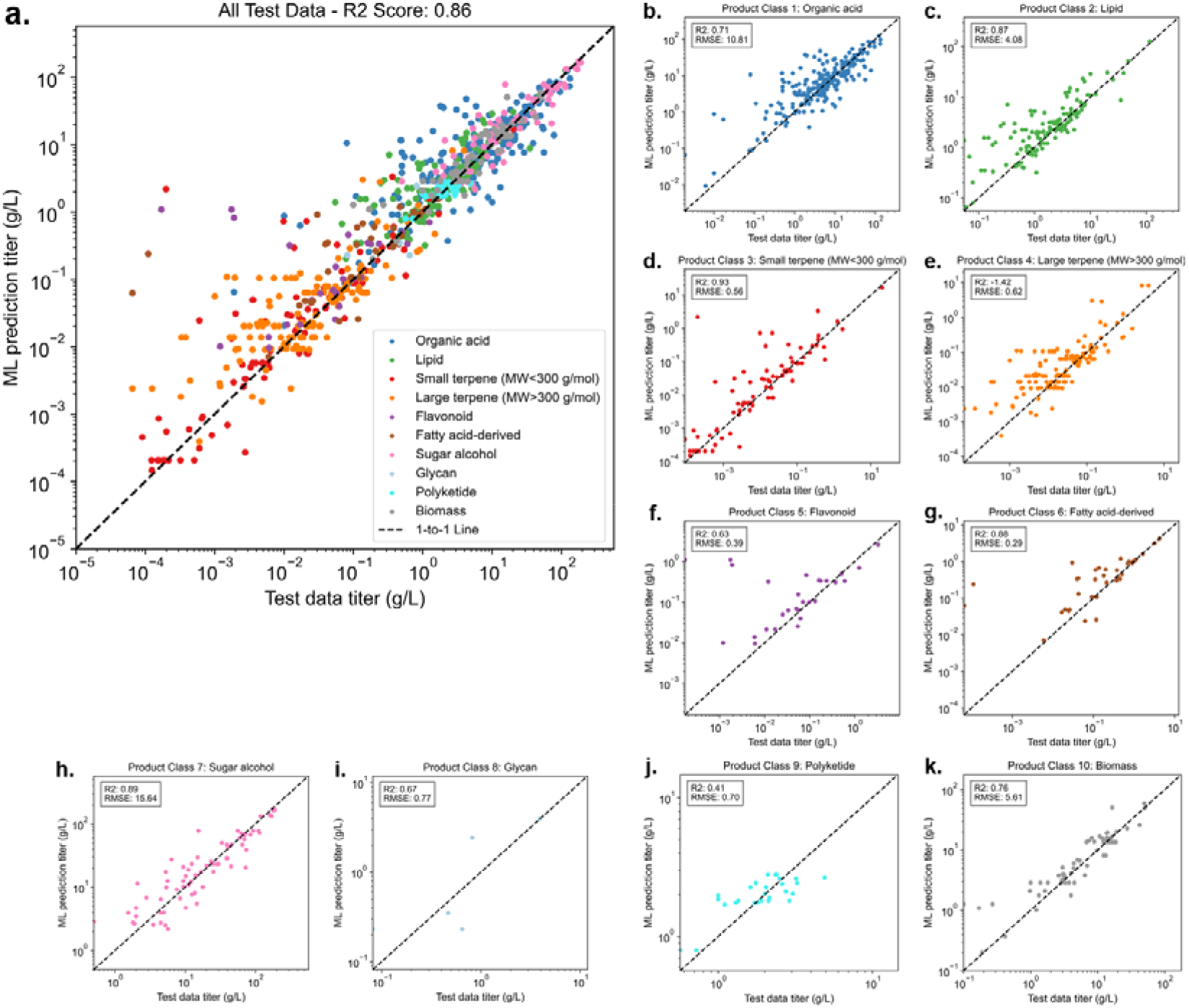
Test set *Y. lipolytica* titer predictions using a random forest ensemble learner.

### Transfer learning from *Y. lipolytica* to *R. toruloides* for prediction of non-model yeast factories

Non-model cell factories have been rapidly developing due to their unique metabolic capabilities. For example, *R. toruloides* is a non-model yeast that can convert cheap feedstock to high-value carotenoids. However, non-model cell factories have a limited number of reports. To better predict their performance under novel conditions, we utilized transfer learning to leverage knowledge from the *Yarrowia* dataset. For example, our *Rhodosporidium* database contained 366 fermentation results extracted from 60 articles that could support RF model to predict lipid and biomass production (R^2^ > 0.4) (**Supplementary Figure 4**), but the database lacks genetic engineering features. For instance, reports on astaxanthin production in *R. toruloides* mainly focused on its native pathways^34–36^. Therefore, knowledge transferring from *Y. lipolytica* papers may predict how genetic engineering affects astaxanthin production in *R. toruloides* and offer guidelines for future strain development. Here, we utilized two inductive learning approaches^37^: 1) a neural network with an pretrained encoder-decoder structure to conduct presentation learning of the effect of the number of gene expressions on the astaxanthin synthesis; 2) an instance-based random forest TL approach to address the source-target domain gap. We evaluated these two different approaches because they have different underlying assumptions. A pretrained encoder implies that *Yarrowia and Rhodosporidium* data are from the same knowledge domain and have the same statistical distribution. In contrast, the instance-based random forest model can handle data from two different knowledge domains.

First, a pretrained encoder in an autoencoder^38^ was used to minimize the effect of noise in the data through feature reduction. This method reduced the number of features from 29 (original 28 features + 1 categorical input of species, *Yarrowia or Rhodosporidium*) to 14. This model predicted the astaxanthin production after 96 hours of growth of *R. toruloides* in shake flasks containing YPD media. The predictions of the model trained on the encoded inputs were unsatisfactory because the model was insensitive to genetic modification features (**Supplementary Figure 5**). Specifically, strains with 0, 2, or 4 heterologous gene overexpression were predicted to have similar astaxanthin titers. Encoding input data can impair the RF’s ability to learn biologically significant knowledge among data, especially genetic modification features. Additionally, the stochastic nature of the encoder’s neural network made this method insensitive to titer differences at the milligram level in a small dataset with little variance in features. The average product titer for astaxanthin was 0.00097 g/L, while the average product titer for all product types for *Y. lipolytica* and *R. toruloides* were 9.3 g/L and 6.4 g/L respectively. Therefore, we concluded that feature dimension reduction by encoding data in TL may have a detrimental effect when applied to complex and interconnected biological systems.

As an alternative to the encoder approach, a RF transfer learning method was tested. The rationale for using a RF is that this algorithm has been shown to make accurate predictions from preliminary tests. During model training, the training instances were labeled as either *Yarrowia* or *Rhodosporidium* (categorical input feature), and a weight of 3× was assigned to the *Rhodosporidium* data. The model trained on data from both species was used to make *R. toruloides* astaxanthin titer predictions. Again, the inputs used corresponded to a 96-hour fermentation in shake flasks containing YPD media. The model predicted that wildtype *R. toruloides*, without heterologous genes, would produce an average astaxanthin titer of 4.2 mg/L, with most predictions below 2 mg/L (**Figure 6**). This is comparable with a recent publication (not used in the model training) reporting astaxanthin production of 1.26 mg/L by *R. toruloides* in shake flasks^36^. Increasing the number of expressed genes from 0 to 6 resulted in increased titer predictions, but with diminishing returns for each additional heterologous gene. The highest titer predictions, an average titer of 39.5 mg/L, corresponded to over expressions of six heterologous genes. The broad distribution of astaxanthin production observed in the bootstrapping analysis suggests that genetic variations beyond the number of expressed genes play a crucial role in determining potential astaxanthin titer. For *Y. lipolytica*, high astaxanthin titer was typically achieved by employing heterologous enzymes with high activities^39, 40^. However, our current model is unable to account for molecular-level differences between enzymes of the same biosynthesis reaction. This RF transfer learning attempt is in essence a summary of strain engineering experiences in *Y. lipolytica*, with transferred knowledge from the growth characteristics of *R. toruloides*.

**Figure 6.**
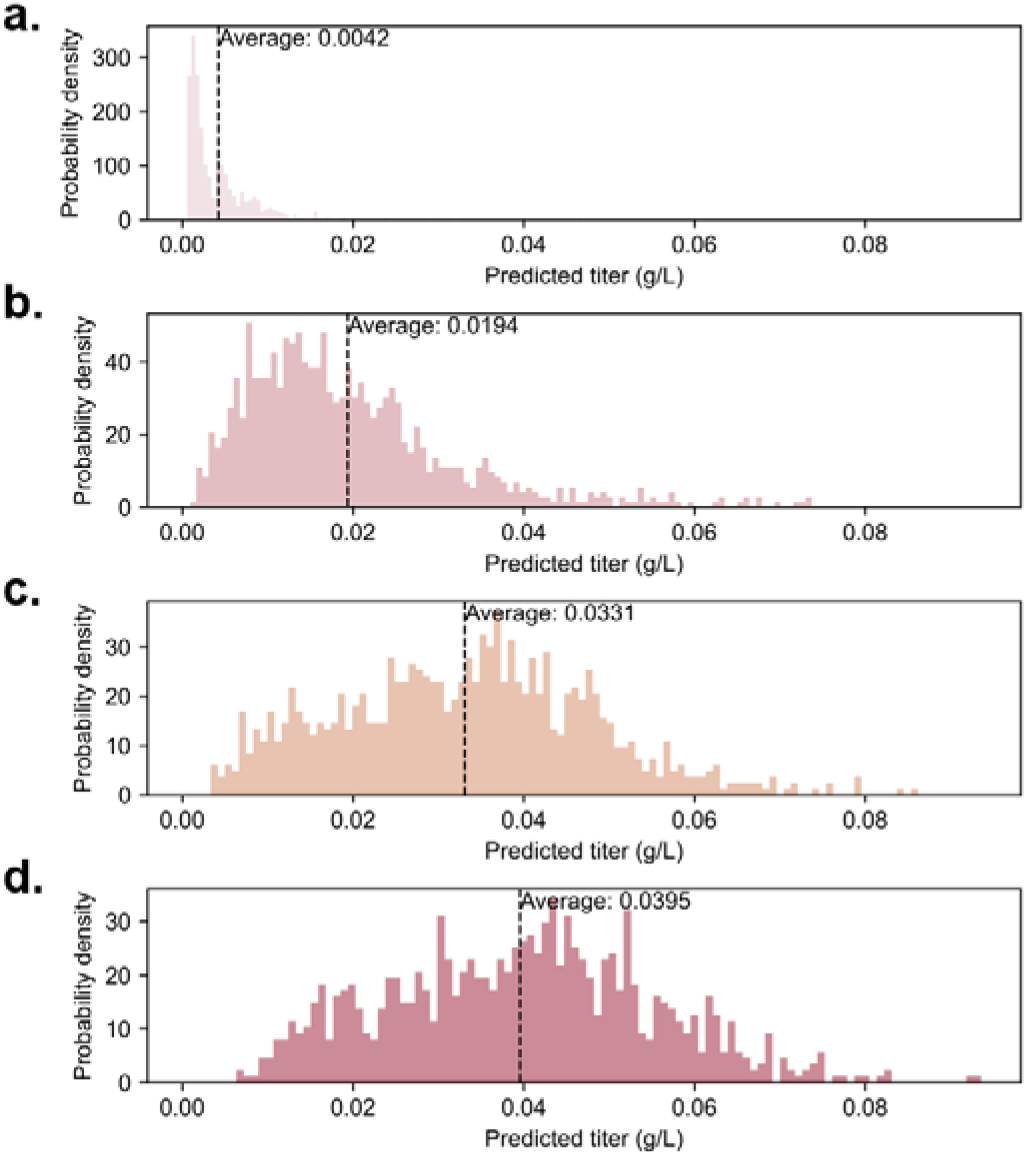
TL prediction for astaxanthin production in *R. toruloides*. **a** 0 heterologous gene expressed. **b** 2 heterologous genes expressed. **c** 4 heterologous genes expressed. **d** 6 heterologous genes expressed.

### GPT-4’s limitations, tips, and future ML/NLP direction

This study also found the limitations for using GPT-4. First, we experimented with GPT-4’ ability to integrate data tables and output the final ML-ready dataset, and it showed satisfactory performance. However, graphical data are still excluded from GPT-4’s data extraction process. In the future, multimodal language models can interpret images for data extraction^15, 41^. Second, we experimented with GPT-4’s ability to integrate data tables and output the final ML-ready dataset, and it showed satisfactory performance. However, this process would stop after a few rows due to token limitations (around 8k), and context length is a bottleneck for large-scale GPT applications. Third, during our data extraction process, we encountered instances when GPT-4 was unable to differentiate between promoters and genes, or mixed up native and heterologous genes. Without knowledge about GPT-4’s self-supervised learning mechanism, it is difficult to explain its performance because of its non-deterministic nature. Fourth, GPT occasionally provides seemingly plausible yet non-factual answers due to misunderstanding the prompt. To resolve these problems, techniques such as few-shot prompt engineering, Chain of Thought, augmented language models, or reinforcement learning through human feedback can be applied ^15, 42^. Fifth, GPT-4 struggles to make good predictions beyond its database. Transfer learning is the solution when researching for the unknowns. With current features, a simple RF with instance-transfer method showed a good ability to give reasonable generalization predictions on unseen data when the size of database is not large. For microbial hosts, such as *S. cerevisiae* and *E. coli*, holistic knowledge mining and large databases could be realized by extracting data from tens of thousands of relevant articles.

In the future, the releases of developer tools and plugins of GPT-4 have the potential to revolutionize data science in synthetic biology. AI can facilitate rapid data sensing, processing, and the implementation of classical machine learning algorithms to accelerate the DBTL cycle.

### Concluding remarks

Although artificial general intelligence is still in its infancy, it has shown great potential to significantly assist and even revolutionize basic scientific research. Our work shows that it is feasible to leverage the power of GPT to automate knowledge mining from existing literature for supporting ML applications. AI models like GPT-4 can process vast amounts of information, reducing the effort researchers need to spend on literature analysis. We also show that this ability, combined with transfer learning, can enable us to discover new knowledge on domains with limited data. We anticipate that generative AI will have broad applications in biological research, including computational strain design, fermentation optimization, prediction of microbial production outcomes for techno-economic analyses, and automation of literature reviews and data interpretation.

## Methods

### GPT-3.5 and GPT-4 versions

The GPT-3.5 version^42^ used was between the dates 3/10 to 3/15/2023. The GPT-4 version was between 3/16 to 4/2/2023.

### Online database

The extracted datasets were deposited in ImpactDB (https://impact-database.com/). This database is currently under development as we are ramping up to upload more data and adding more online tools.

### Data extraction from journal articles

Data extraction and feature organizations (**Supplementary Figure 1**) were done in a semi-automated fashion because currently GPT-4 is only available through OpenAI’s ChatGPT website. First, text files for the different article sections (title, abstract, method, results, and text-based tables) were generated by manually combining the prompt sentence from **Table 1** with corresponding text that was copied-and-pasted from a manuscript. The prompt sentences were designed so that the extracted data from this study has the same format as the data from previous *Yarrowia* knowledge engineering effort^2^. GPT-4’s responses were recorded and then later combined into a ML-ready format. The data correctness test was done by applying the above workflow to 10 articles of the *Yarrowia* paper database in ImpactDB. We excluded a few products not in the top ten product classes from our ML training due to their small sample volume. However, the data related to these products was uploaded to ImpactDB. The extracted data and paper number assignment were reported in **Supplementary File 2**. Standard condition thermodynamics data were obtained from NIST webbook (https://webbook.nist.gov/chemistry/). Non-standard thermodynamics data under biologically relevant concentrations were estimated by eQuilibrator 3.0 (https://equilibrator.weizmann.ac.il/27). Pathway enzyme steps, ATP, and NAD(P)H costs were estimated either by consulting KEGG^43^ (https://www.genome.jp/kegg/) or reading the pathway map in relevant journal articles.

### Manual and AI extracted data analysis, Data preprocessing, and ML

All classical ML algorithms were implemented using the scikit-learn Python package. For the calculation of feature variances in comparing manually extracted data and AI extracted data, features were normalized with their mean values and then the variances were calculated. The feature importance was determined by training a RF regressor and extracting the corresponding weights. The clustering of PCA space was done by K-means unsupervised learning using a cluster number of 6, and Euclidean distances was used to determine the distance between cluster centroids. Data preprocessing for both ML and TL are specified in the following steps. A simple imputation was performed, with missing values as zero to maintain its biological meaning. Stratified data split was performed to ensure that at least 20% data of each product class were included in the testing data. Categorical data are encoded using ordinal encoding, enabling feature comparison after training and testing.

### TL via Pretrained-encoder and RF with Instance-transfer

Transfer learning is a machine learning paradigm that leverages knowledge gained from solving one task and applies it to improve the performance of a related but distinct task. The key idea behind transfer learning is to use the learned representations or knowledge acquired during the training of a pre-existing model, often referred to as the “source” task, and apply this knowledge to a different, “target” task. In this work we mainly adopt two approaches: fine-tuning a pre-trained encoder-decoder and instance transfer on Random Forest. Fine-tuning usually entails continuing the training process with a smaller learning rate on the target task data, allowing the model to update its weights and adapt to the new task while preserving the valuable knowledge gained from the source task, while instance transfer mainly involves combining the source and target dataset. We first trained the autoencoder structure on all *Y. lipolytica* as source data, then transfered the encoder by freezing the layers to re-train it on *R. toruloides* data without the Astaxanthin data. The resulting embedding was then used to test a random forest model. The best result for predicting different numbers of heterologous genes by this method was shown in **Supplemental File 1** using the first model in **Table 3**. The resulting performance metrics for the regression task to predict all Astaxanthin product titer were shown in **Supplemental File 1**. The model architectures were summarized in the following table and illustrated in **Supplementary Figure 6**.

**Table 3.** Details of the encoder-decoder structure.

| Model number | Loss equation | Model | Output dimension |
| --- | --- | --- | --- |
| 1 | $L_{tot} = L_{Reconst} + l1\_weight * L_{l1}$ | 2-layer encoder + 2-layer decoder with normalization | 50% reduction |
| 2 | $L_{tot} = triplet\ loss + l1\_weight * l1\_penalty$ | 2-layer encoder + 2-layer decoder with normalization | 50% reduction |
| 3 | $L_{tot} = L_{Reconst} + l1\_weight * L_{l1}$ | 2-layer encoder + 2-layer decoder with normalization | 0% reduction |

The details of encoder-decoder TL method were tabulated in **Table 3**. Note that all layers were fully connected, and early stopping was applied in both pretraining and retraining to prevent overfitting due to noise and outliers. The loss equation, a quality measure of the model, was tabulated in **Table 3**. Reconstruction loss was calculated as the MSE between the original x and embedded x, and the l1 loss was the regression score calculated after encoding. Triplet loss was calculated as the Euclidean distances between the negative sample/positive sample and the anchor then applying the Rectified Linear Unit (ReLU) on the difference between the distances added by the margin. Autoencoder’s neural network weight was initialized using He initialization^38^ for all experiments. The corresponding 0, 2, 4 heterologous gene expressions samples were encoded as input data and the prediction was performed by training a separate RF on a synthetic dataset obtained by bagging with replacement 100 times on all Astaxanthin data since the sample size for Astaxanthin is small.

The RF instance-transfer was done by the following steps. To balance the *R. toruloides*’ effect on the prediction due to its small size, we augmented the *R. toruloides* data to three times its initial size to form the combined dataset. The same data preprocessing was followed. The model was then tuned on the new training set and prediction is done using Astaxanthin samples with 0, 2, 4, 6 heterologous genes expressed respectively. The trained model was used to make predictions on the product titer. All codes are available at https://github.com/wenyuli23/SyntheticBiologyTL.

### Statistical methods

All data were reported as the mean ± standard deviation. Statistical test used in this study was a one-tail student t-test.

## Acknowledgements

This study is funded by United States NSF award number 2225809. The authors GR and YJT also received funding from Defense Advanced Research Projects Agency B-SURE program (HR001122S0010) for this research. The views, opinions and/or findings expressed should not be interpreted as representing the official views or policies of the Department of Defense or the U.S. Government.

## Author contributions

Conceptualization and Ideas - YJT, YC and GR; Methodology and Programming – GPT-4, WL and ZX; Data organization, curation, website, and deposition - GR, ZX and HM; Writing – ChatGPT, WL and ZX; Review & Editing - YJT, YC, HM.

## Supplementary files

Supplementary file 1: GPT-4’s response screenshots and additional details of the machine learning workflow.

Supplementary file 2: GPT-4-extracted *Y. lipolytica* bioproduction database

## TOC graphic

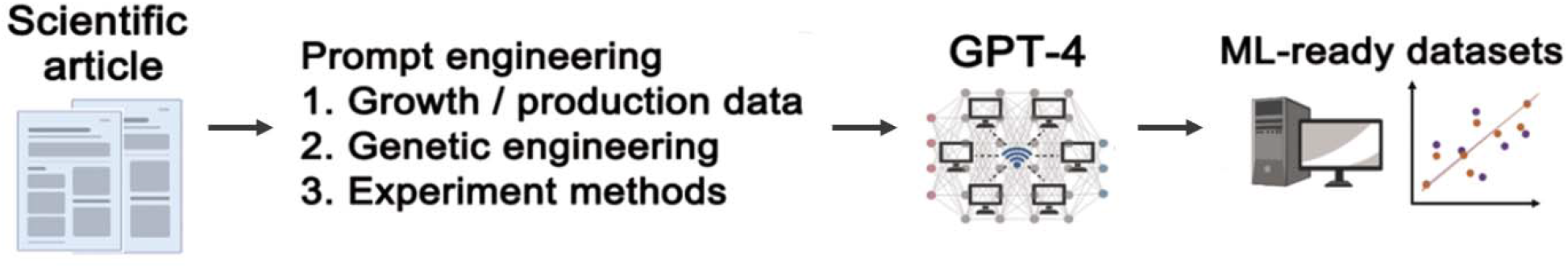

