## Supplemental file 1 for "Generative artificial intelligence GPT-4 accelerates knowledge mining and machine learning for synthetic biology"

### Supplementary file 1

**Generative Artificial Intelligence GPT-4 Accelerates Knowledge Mining and Machine Learning for Synthetic Biology**

Zhengyang Xiao^a,§^, Wenyu Li^b,§^, Hannah Moon^c,d^, Garrett W. Roell^a,b,c,*^, Yixin Chen^b,*^, Yinjie J. Tang^a,*^

^a^ Department of Chemical Engineering, Washington University in St. Louis, St. Louis, MO, 63130, USA

^b^ Department of Computer Science and Engineering, Washington University in St. Louis, St. Louis, MO, 63130, USA

^c^ ImpactDB LLC. St. Louis, MO, 63105, USA

^d^ Clayton High School, 1 Mark Twain Cir, Clayton, MO 63105, USA

^§^: These authors contributed equally to this work

***Corresponding Authors:**

Garrett Roell ([](http://))

Yixin Chen

Yinjie Tang

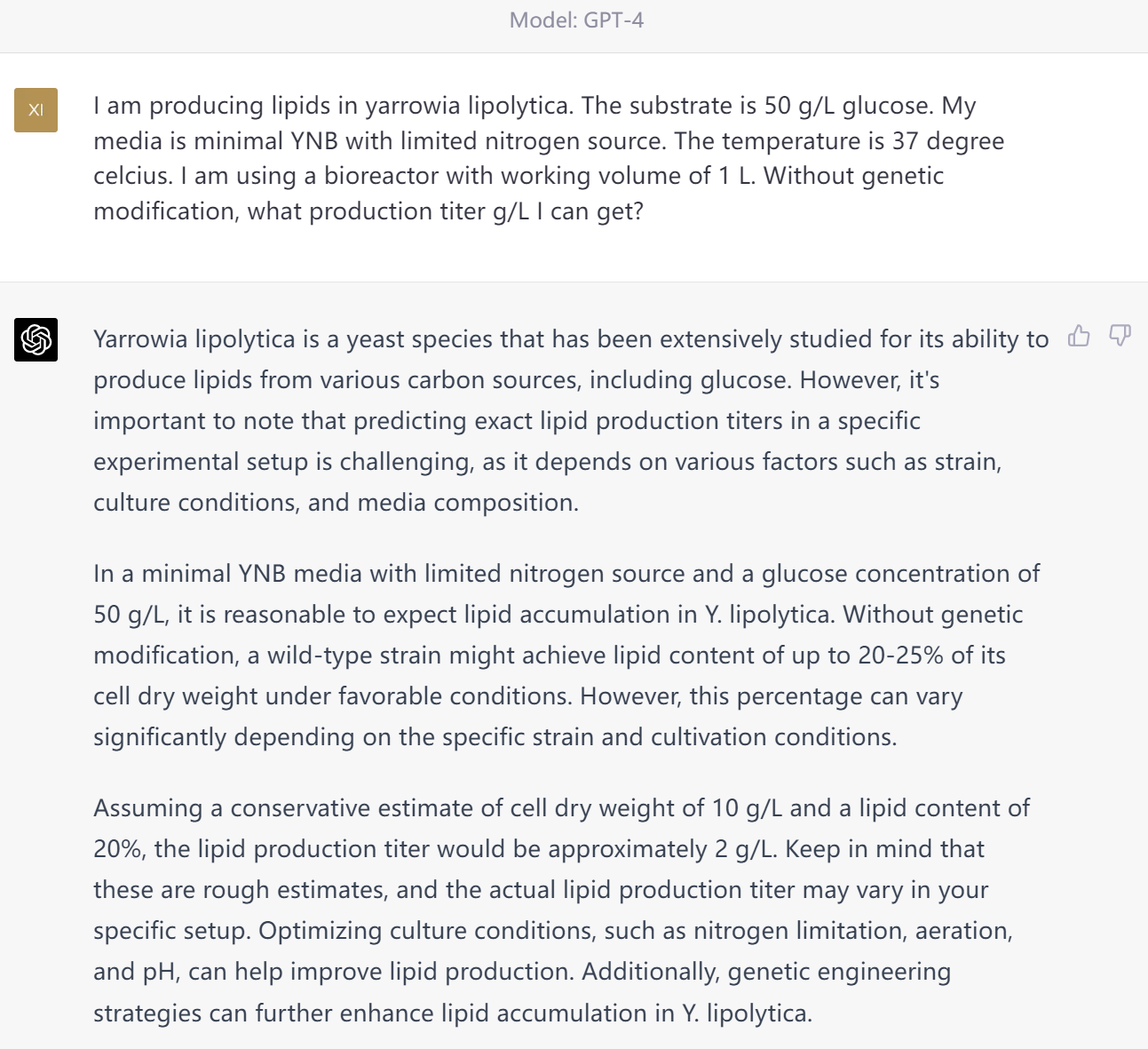

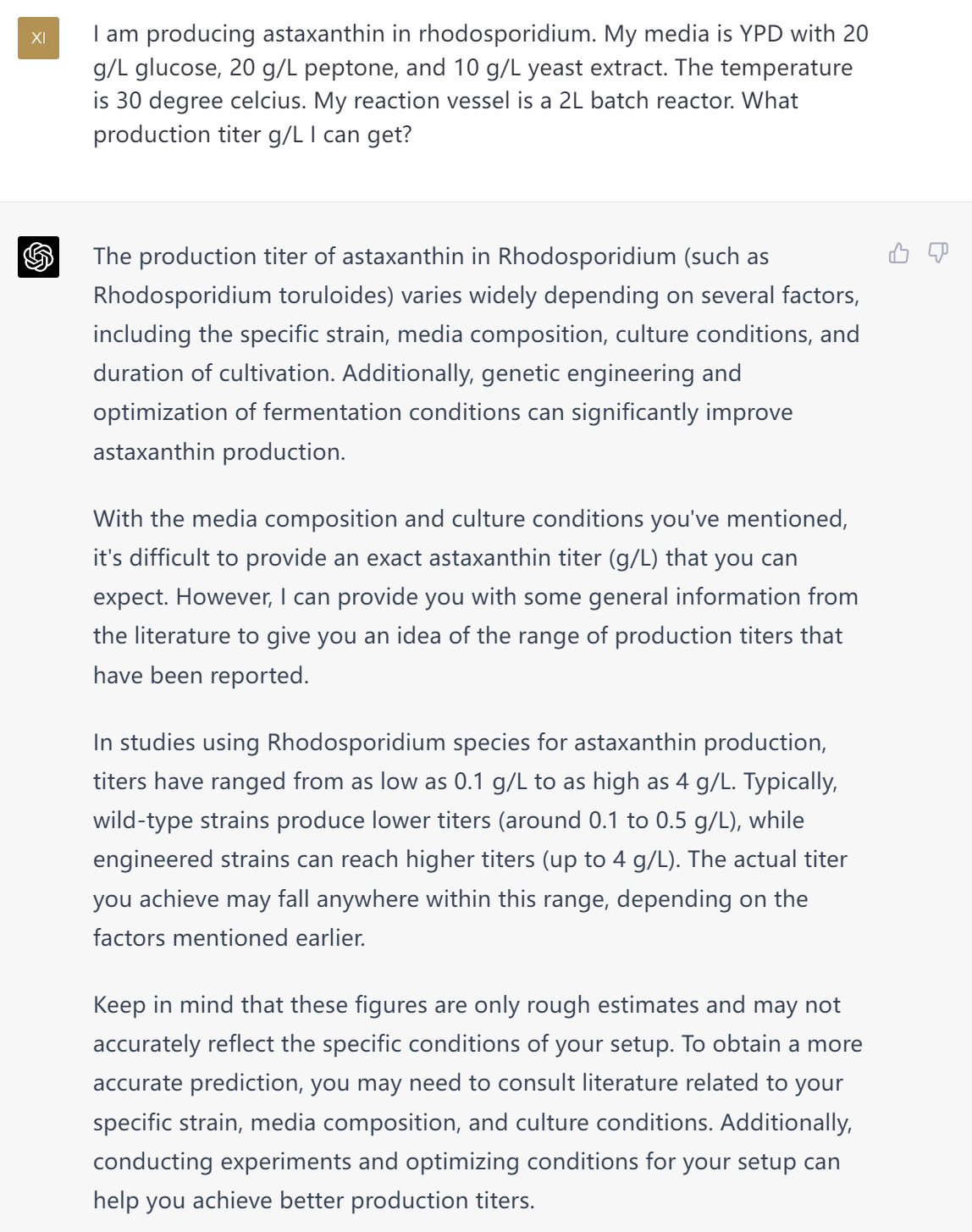

**Supplementary Figure 1. GPT-4’s estimate of lipid production in *Yarrowia lipolytica* and astaxanthin production in non-model yeast *Rhodosporidium*.**

The oleaginous yeast *Y. lipolytica* typically accumulates lipids more than 40% of its dry cell weight under nitrogen limited conditions. The biomass yield from glucose is around 0.3 g/g. A well-trained microbiologist would give a lipid titer estimation of 0.3*50*0.4 = 6 g/L, which is comparable to GPT-4’s prediction. The lipid production is dependent on several environmental factors, as our machine learning model has gram-level errors when predicting lipid production titer. However, GPT-4 ignored an important factor, that is *Y. lipolytica* cannot grow under 37°C.

Very few research papers have been published on astaxanthin production in *rhodosporidium*, with the highest reported titer 0.0013 g/L in Tran et al.^1^ in the year 2023 (GPT-4’s database was up to 2021). Under the given conditions, GPT-4’s estimate appeared to be unrealistic.

**Supplementary Table 1 Comparison of manual data extraction with GPT-3.5 and GPT-4**

|  | **Manual** | **GPT-3.5** | **GPT-4** |
| --- | --- | --- | --- |
| Source data location | Text, figures, and tables | Text and tables | Text and tables |
| Output data format | Manually typed spreadsheets | Data tables in markdown format (copied into spreadsheets) | Data tables in markdown (copied into spreadsheets) |
| Correctness of data | Eyeballing figures can create errors. Data from text and tables are exact.  Variation in correctness between human readers | Some are wrong. Others are exact. Test set accuracy = 89% | All data are exact. Test set accuracy = 100% |
| Correctness of experiment methods | Mostly good. | Mostly good, sometimes failed to give a table. | Good, GPT-4 can reason through the context (Figure captions, discussions, etc.). |

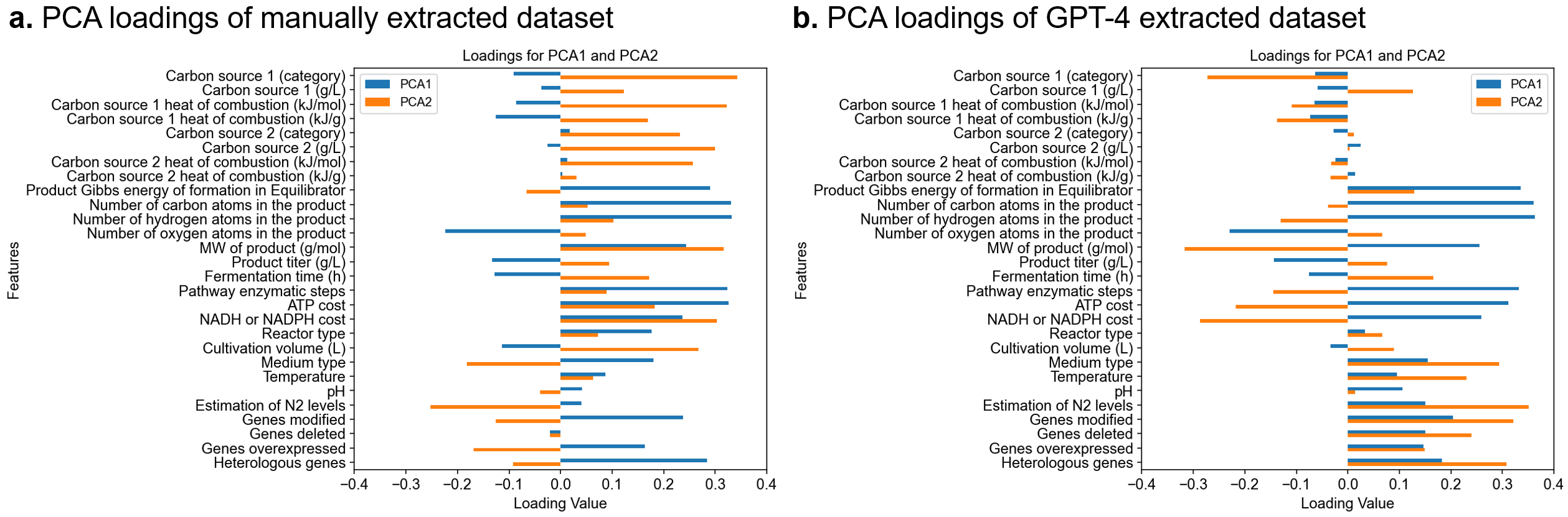

**Supplementary Figure 2. PCA loadings of two datasets. a. PCA loadings of manually extracted dataset.** Most loadings concentrate in carbon source and production pathway costs. **b. PCA loadings of GPT-4 extracted dataset.** Most loadings concentrate in growth condition, genetic engineering, and production pathway costs.

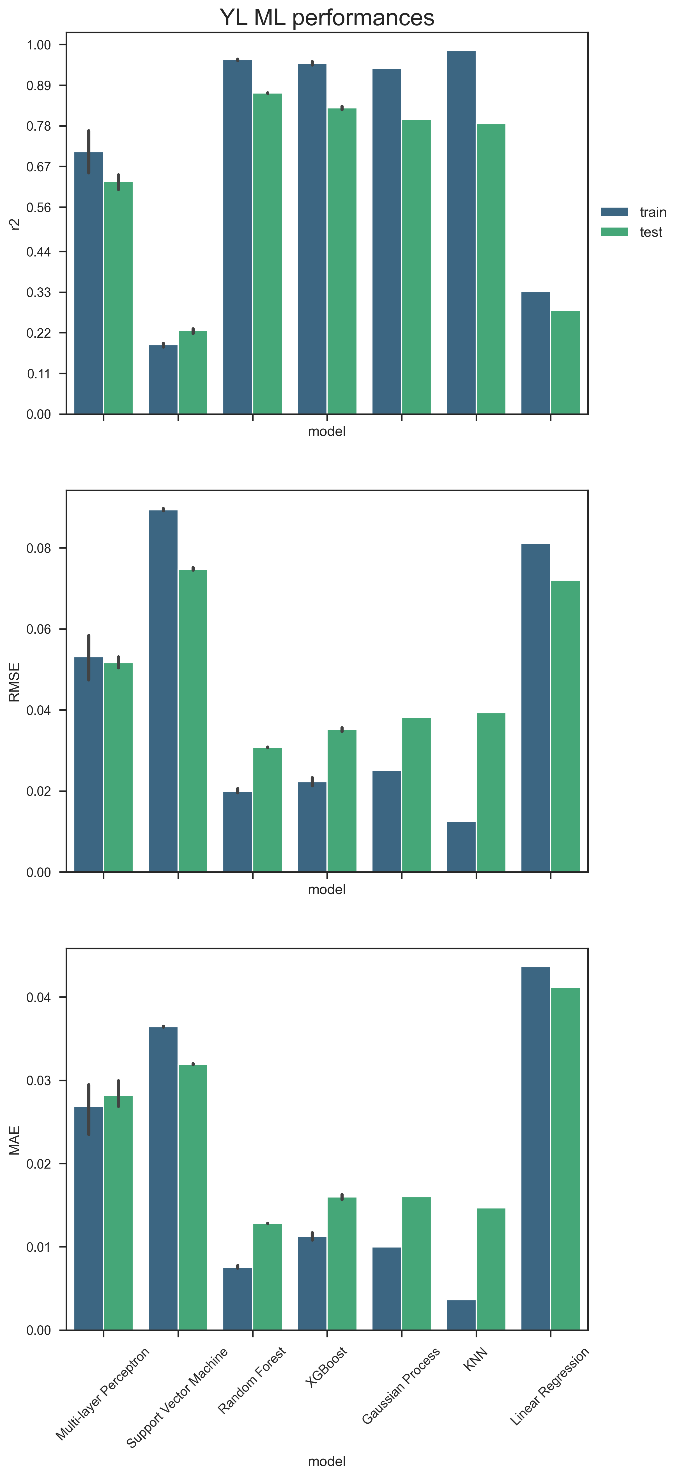

**Supplementary Figure 3. Comparison of ML performance when predicting training and testing instances for *Y. lipolytica*.** The y-axis is the value of the metric, and the x-axis is the different algorithms, with the bar value as the mean, and the black line as the standard deviation. The variation in values is obtained by optimizing RMSE (root mean square error), MSE (mean square error), R^2^, and MAPE (mean average % error). The titer data were scaled when comparing different ML algorithms.

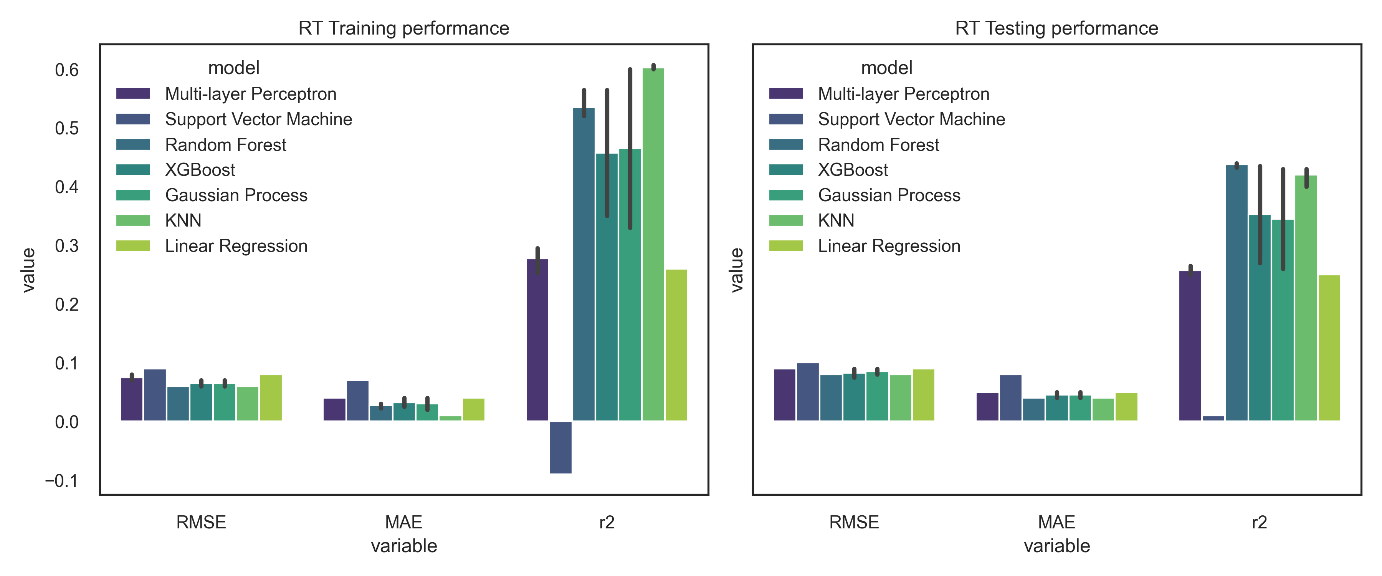

**Supplementary Figure 4.** **ML model training and testing performances for *R. Toruloides*.** With the bar value as mean, the black line as standard deviation.

**Supplementary Table 2. Parameters chosen for ML models for YL**

| MLP | 2-layer structure, with rectified linear (ReLU) activation function and early stopping applied |
| --- | --- |
| SVM | Linear kernel |
| RF | MSE with Friedman’s improvement is used for splitting criterion, and bootstrapping is used |
| XGBoost | gamma = 0, learning_rate = 0.2, max_depth = 5 |
| KNN | Manhattan distance |
| GP | a RBF kernel with length scale 0.5 added by a white kernel with noise level 0.1 is used |

a

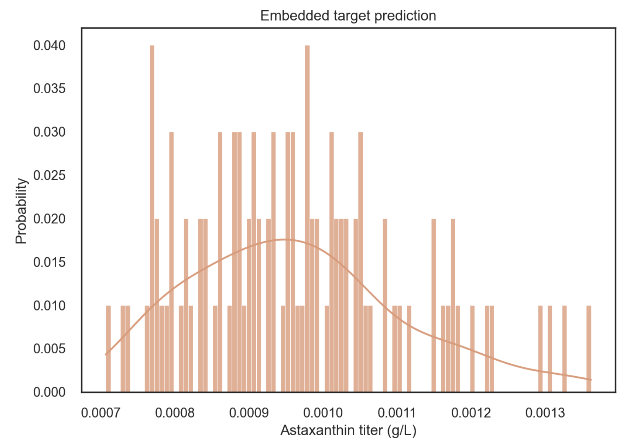

b

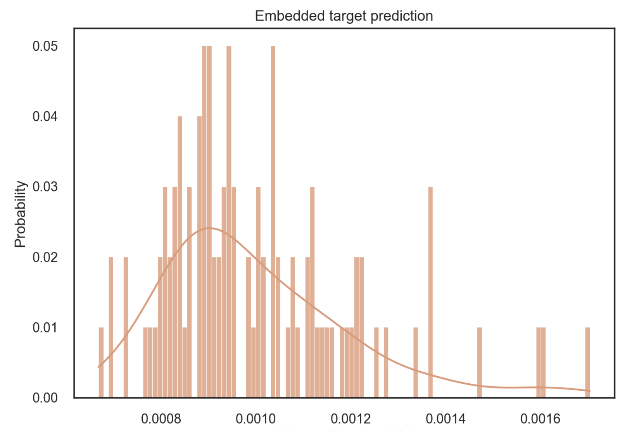

c

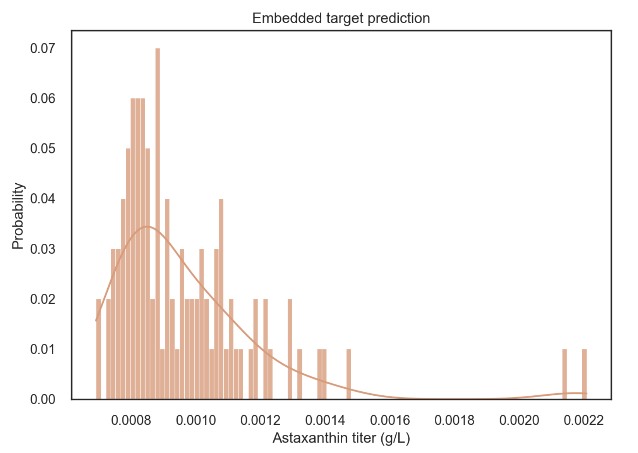

**Supplementary Figure 5. Autoencoder transfer learning prediction on astaxanthin production in *R. toruloides*. a** 0 heterologous gene expressed. **b** 2 heterologous gene expressed. **c** 4 heterologous gene expressed.

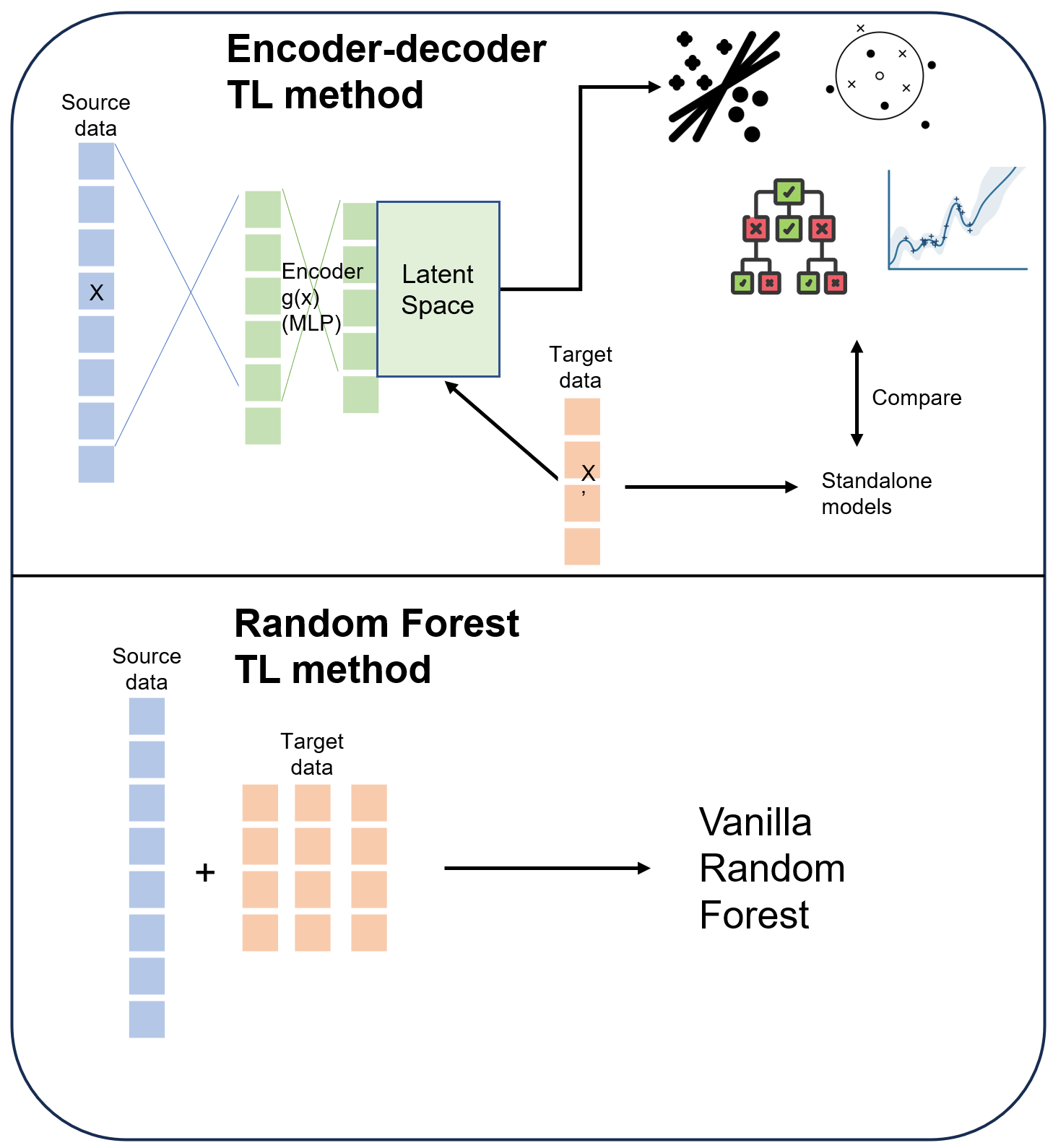

**Supplementary Figure 6. Graphical illustration of transfer learning model structure.**

**Screenshots of GPT-4 responses in our AI-enhanced workflow**

Below is GPT-4’s response to prompt 1 (growth condition and thermodynamics data). The text is from a study producing flavor lactones in *yarrowia*^2^. The prompt was added before the selected text. GPT-4 identified 10 experimental data instances in the response. Although the response is not perfect, it provides a framework for *targeted* search for experiment data, without the need to read all the text.

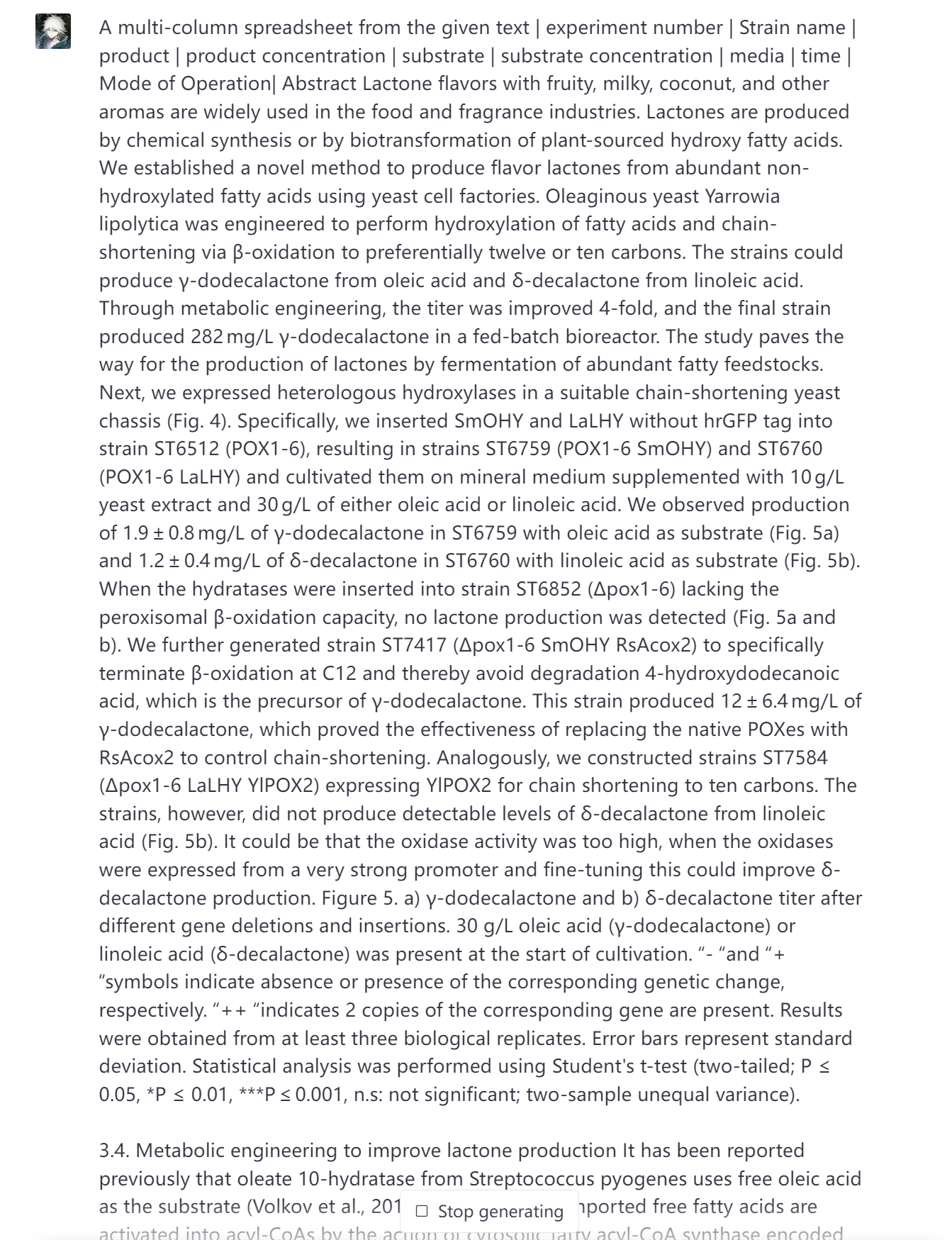

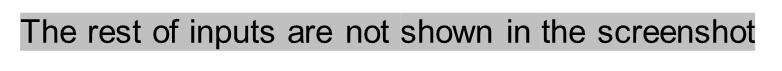

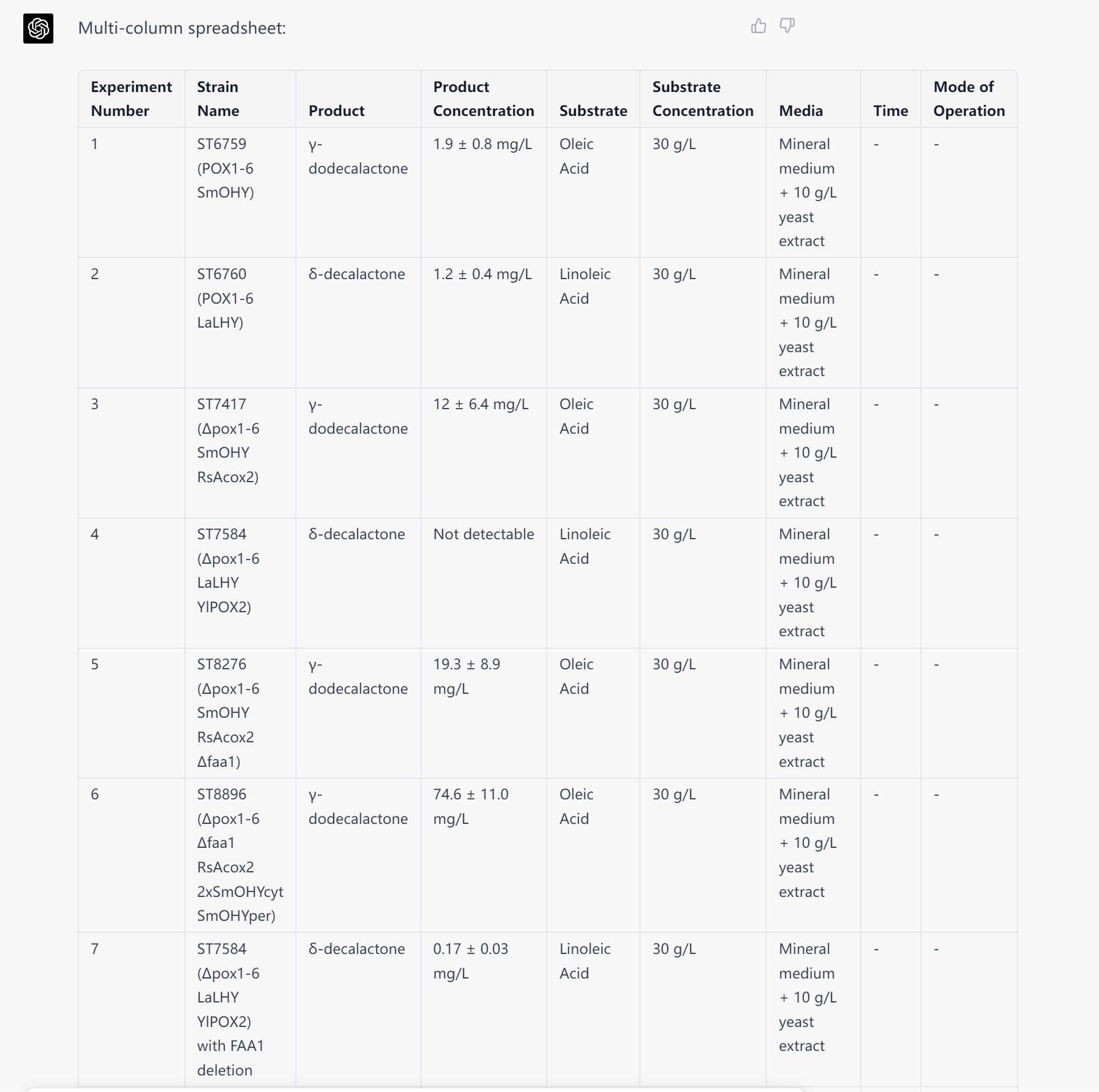

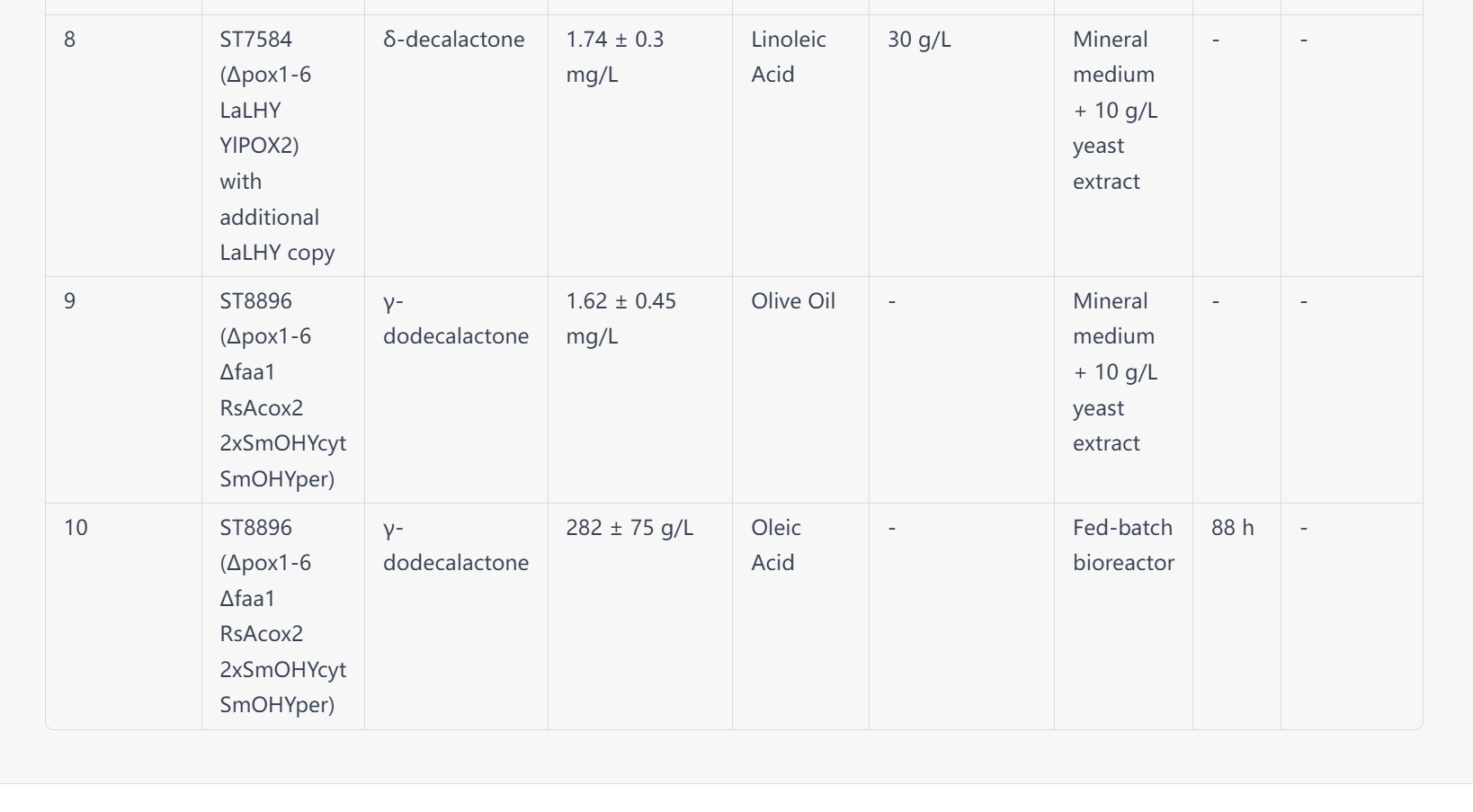

Below is a sample response to prompt 2 (genetic engineering). The text is from a study producing polyols in *yarrowia*^3^. The prompt asks GPT-4 to summarize the strain engineering features including strain name, parent strain, knockout genes, expressed genes, promoter, genome integration, and codon optimization. Since GPT-4 can reason through context, we ask it to include the gene expressed and knockout in the parent strain when outputting the daughter strain.

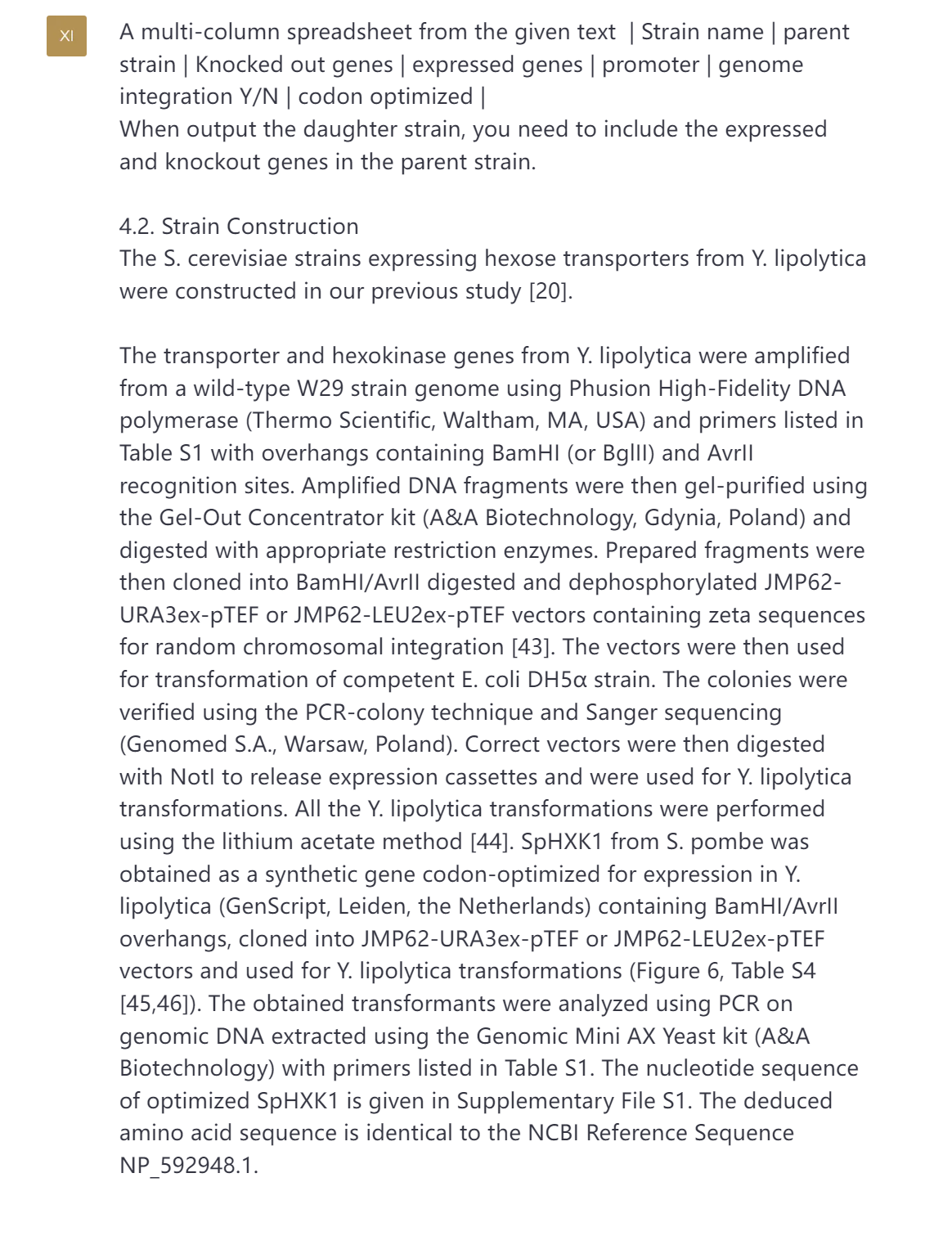

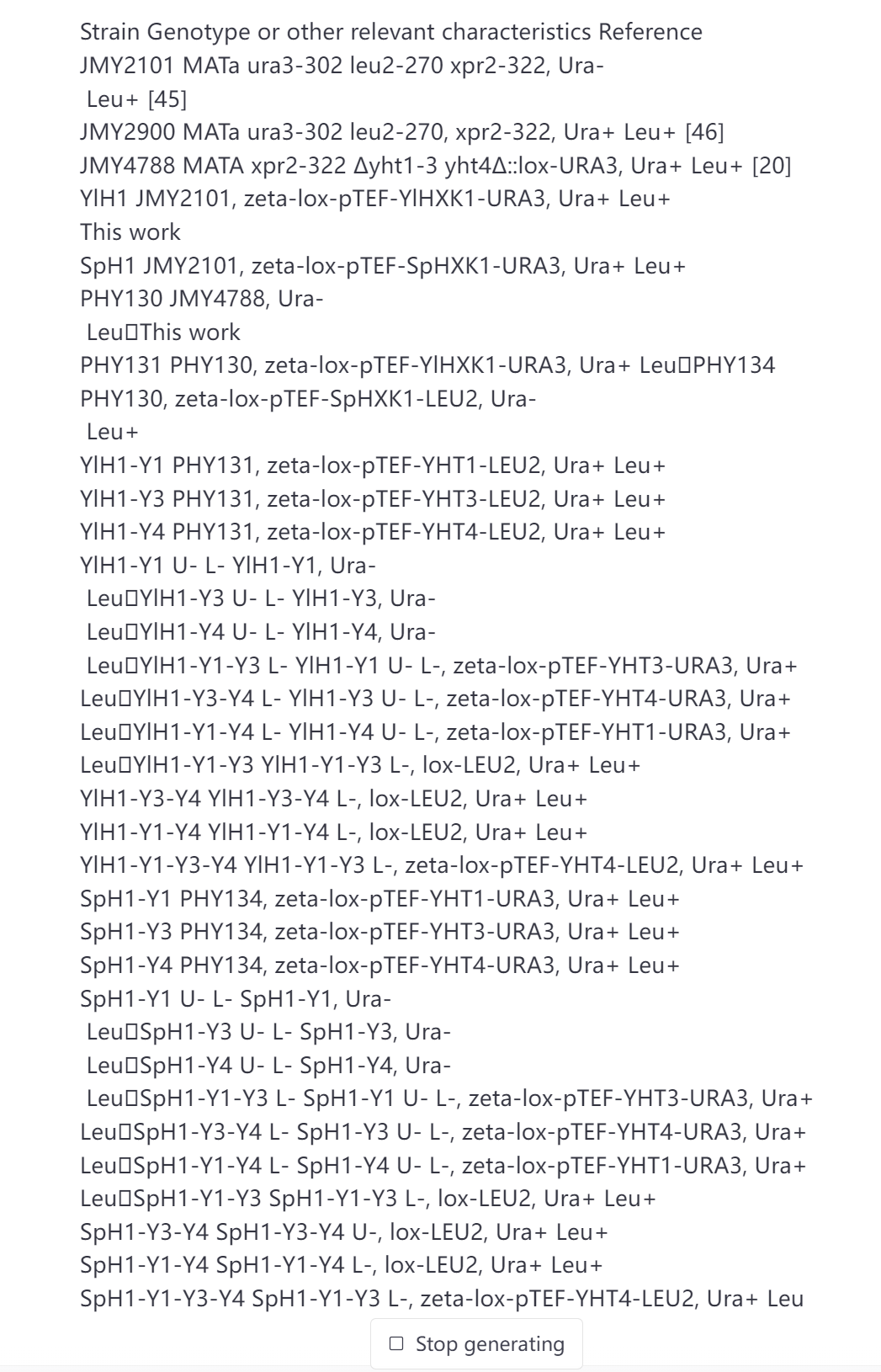

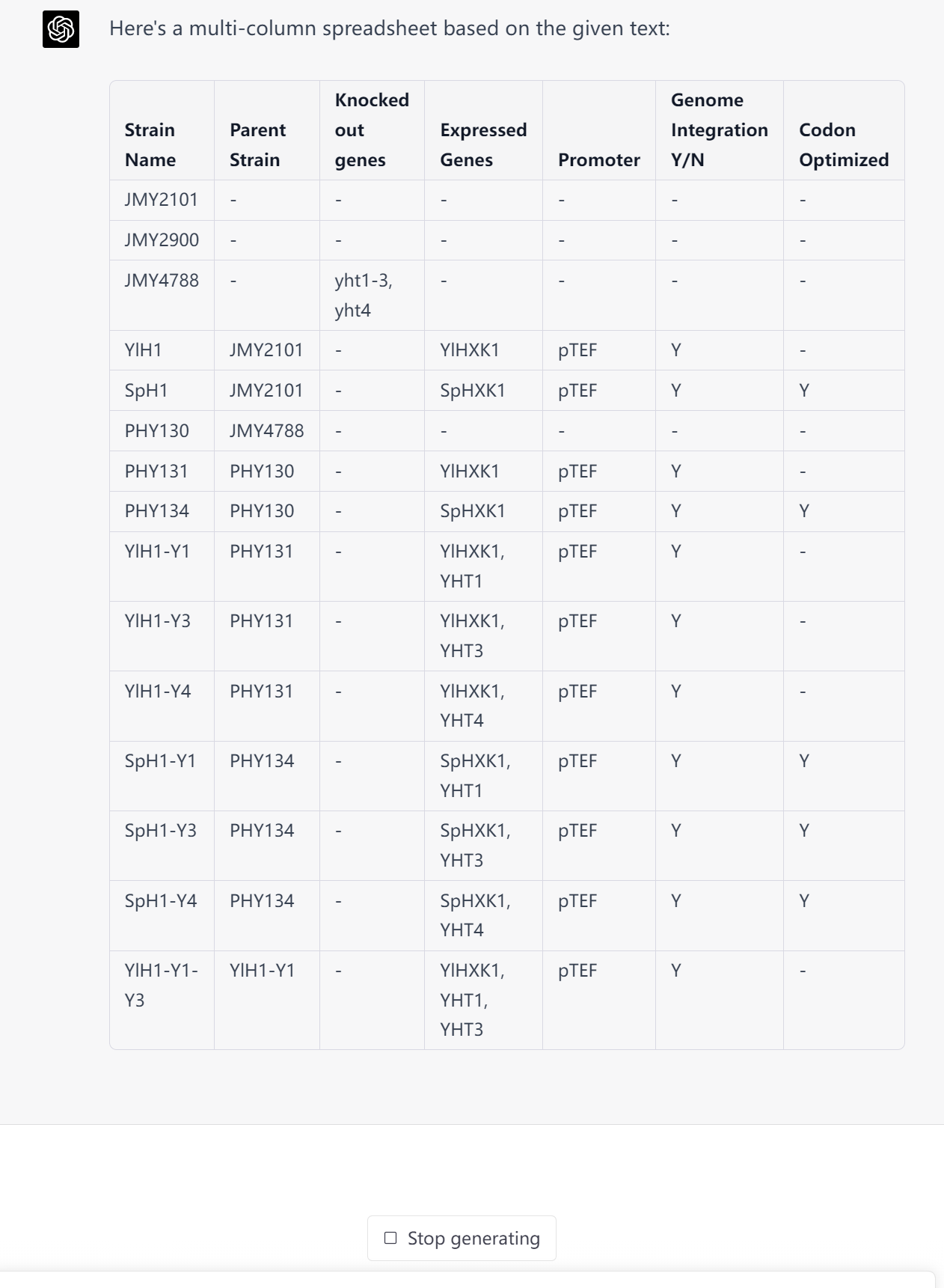

Below is a sample response of prompt 3 (Growth condition). The text is from producing flavor lactones in *yarrowia*^2^. The prompt words were put before the method section. GPT-4 identified 4 types of experiments and their methods.

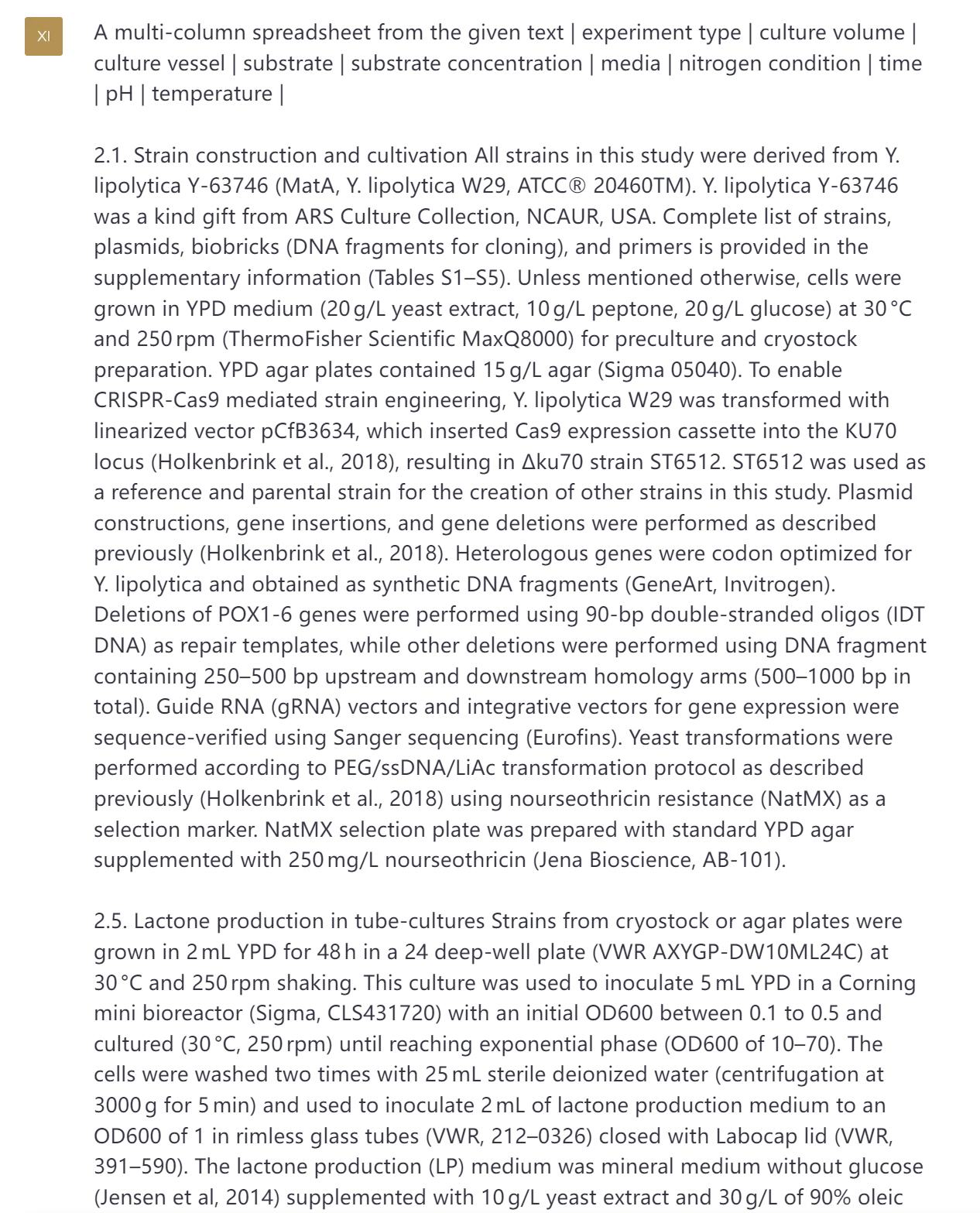

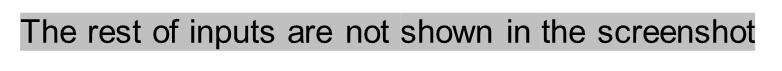

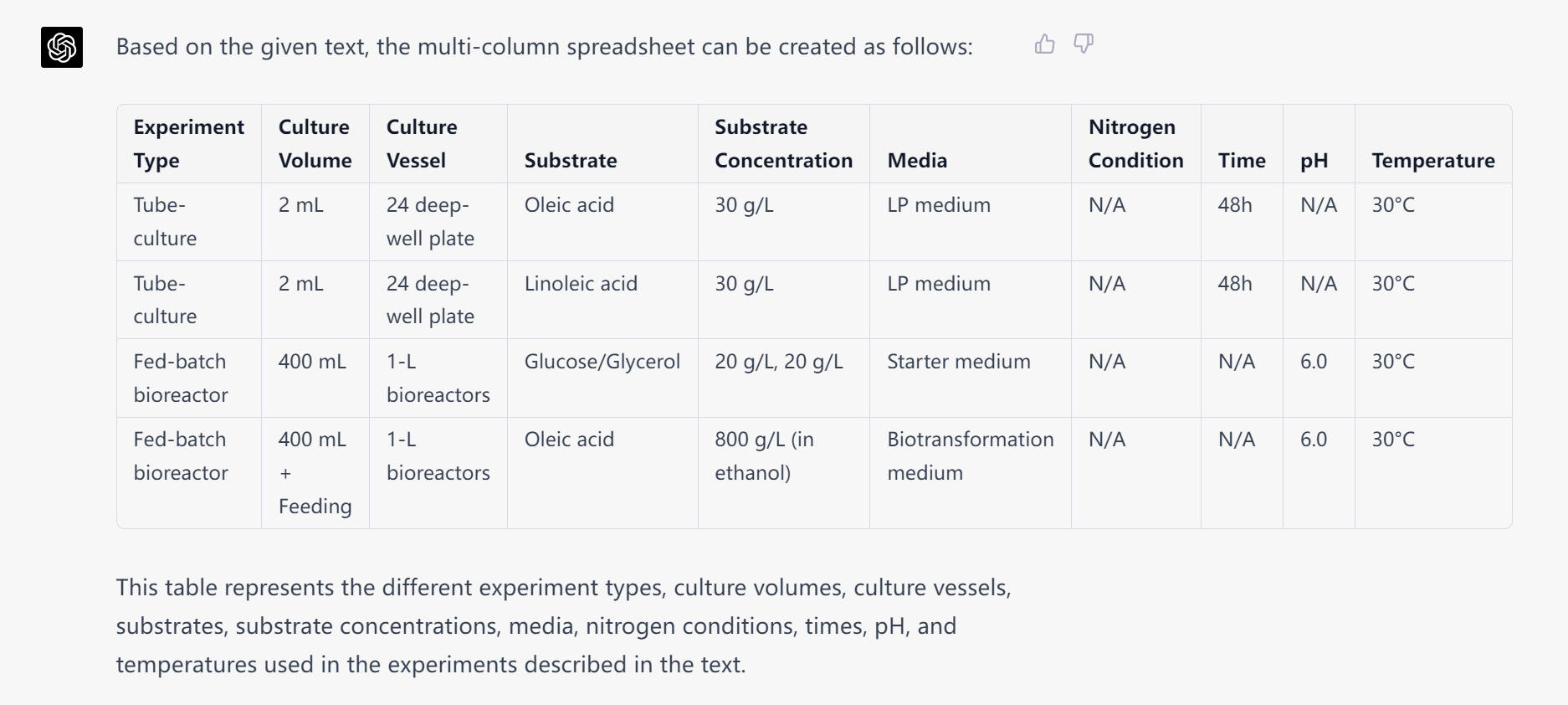
